# Early-life Exposure to Arsenic Primes the Offspring to Increased Asthma Risk: Transcriptome and Epigenome Analysis

**DOI:** 10.1101/2025.08.01.668223

**Authors:** Yuchen Sun, Musa Watfa, Qin-ying Sun, Dhivyaa Rajasundaram, Ben Schlegel, Hoi-yee. Yeung, Jarius Pulczinski, Bongsoo Park, Aaron Barchowsky, Wayne Mitzner, TaRGET II Consortium, Shyam Biswal, Wan-yee Tang

## Abstract

Inorganic arsenic (iAs) in drinking water is a global health concern. This study tests whether maternal exposure to iAs in drinking water at the WHO provisional level (10µg/L) increases offspring asthma risk via epigenetic reprogramming. F1 mice prenatally exposed to iAs were analyzed at 5 months for blood transcriptome and methylome changes and challenged with house allergens before lung function testing. Prenatal iAs exposure led to increased airway hyperresponsiveness (AHR) and altered inflammation gene expression and DNA methylation changes. Notably, *miR-101c* was epigenetically reprogrammed early in development, with persistent downregulation in both target (fetal and adult lungs) and surrogate (amniotic fluid and blood) tissues. These changes correlated with increased allergic AHR and TGFβ pathway dysregulation. Findings suggest that maternal iAs exposure primes offspring for asthma risk through epigenetic alterations and may inform risk assessment and biomarker development in affected communities.

**KEY FINDINGS:**

1. *In utero* exposure to 10 part per billion (or 10 µg/L, the current WHO and EPA provisional level in drinking water) inorganic arsenic (iAs) increases offspring asthma risk. These results raise concerns about the current safety thresholds for iAs in drinking water (Fig. 1).
2. Transcriptomic and methylome analyses of blood leukocytes from 5-month-old F1 mice revealed that maternal iAs exposure induces transcriptional changes in genes related to allergic airway responses. Pathway analysis highlighted the involvement of *miR-101c* and its connection with TGFβ downstream targets in regulating extracellular matrix signaling, embryonic development, and inflammatory (Figs. 2 and 3).
3. Transcriptional changes in c*ol3a1* and *miR-101c* in blood were strongly correlated with allergen-induced airway hyperresponsiveness (AHR), with similar alterations detected in plasma samples. These findings provide new insights into respiratory health in affected communities and support the development of biomarkers for iAs risk assessment (Figs. 4 and 5).
4. Several gene-specific epigenetic alterations induced by early-life iAs exposure were consistently observed in both surrogate (blood) and target (lung) tissues across developmental stages, offering new panels of easily accessible markers for early detection and monitoring of lung disease risk in offspring. The persistence of these markers over time makes them valuable for predictive modeling and life course studies. (Fig. 6).
5. Downregulation of *miR-101c* was validated in fetal lung and amniotic fluid of the iAs-exposed group, suggesting that epigenetic reprogramming of *miR-101c* is initiated early in gestation (Fig. 7). These findings help uncover causal pathways linking environmental exposures to asthma pathogenesis.
6. Distinct sex-specific patterns in blood transcriptome and methylome alterations in respond to early-life iAs exposure underscore the importance of considering sex as a biological variable in omics research.

**GRAPHIC ABSTRACT:** 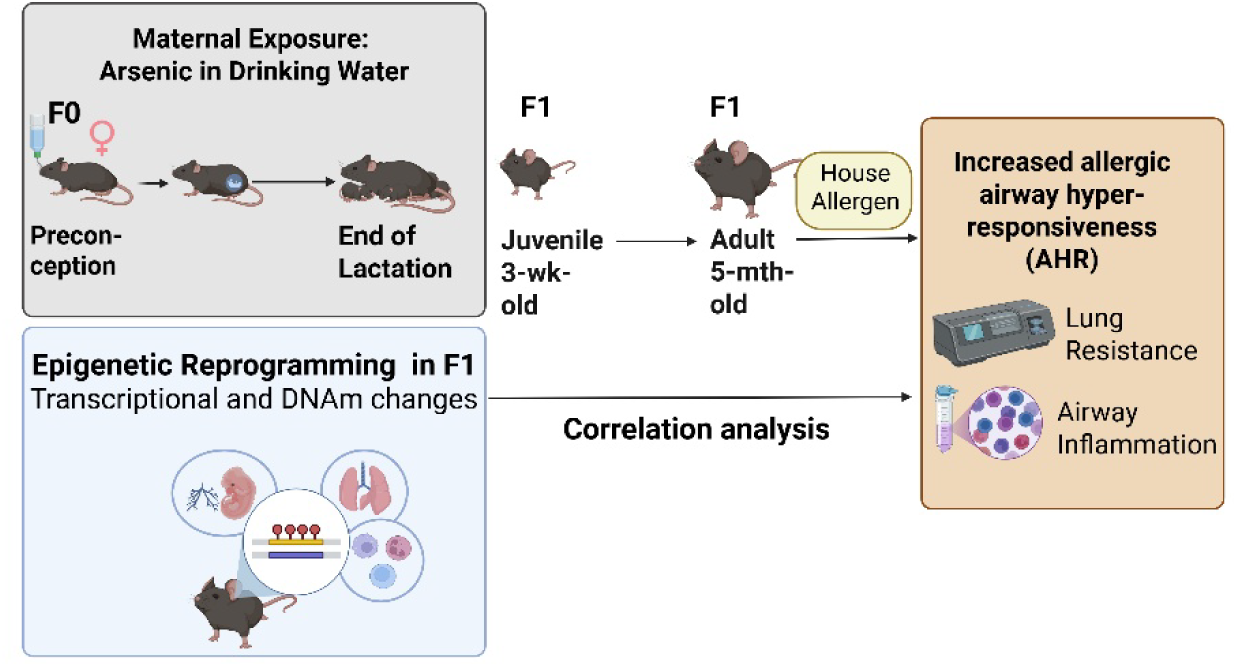

**Description:** F1 progeny prenatally exposed to iAs were assessed for blood transcriptomic and DNA methylome analysis at 5 months of age. Correlation analysis between transcriptional changes and allergen-induced airway hyperresponsiveness (AHR) was conducted to examine the epigenetic impact of maternal iAs exposure on offspring asthma risk.

## INTRODUCTION

Inorganic arsenic (iAs) in drinking water is a major global health concern^1^, with over 200 million people worldwide estimated to be exposed to levels exceeding the WHO and U.S. EPA guideline of 10 part per billion (ppb or µg/L)^2^. In the U.S., 2.1 million individuals rely on private wells containing iAs concentration above this limit^3^. Lifelong exposure to iAs has been linked to respiratory diseases^4^, potentially through impaired immune responses and reduced lung function^5^. Maternal iAs exposure is associated with increased risk of lung inflammation and airway allergy in children^6 7^. Notably, *in utero* iAs exposure adversely affects adult lung function even in the absence of continued postnatal exposure, suggesting the lasting impact of early-life exposures^8^. However, the mechanisms by which early-life iAs exposure contribute to respiratory disease, including asthma, remain poorly understood.

Emerging evidence from epidemiological and experimental studies suggests that low-to-moderate prenatal iAs exposure can induce epigenetic alterations that increase susceptibility to disease in childhood and beyond^9 10^. Both iAs and its methylated metabolites, monomethylarsonic acid (MMA) and dimethylarsinic acid (DMA), can cross the placenta and reach the developing fetus^11 12^. While high-level iAs metabolism may deplete S-adenosyl-L-methionine (SAM), a key methyl donor, such stoichiometric depletion is unlikely at lower exposures. Instead, iAs is postulated to disrupt epigenetic regulation of gene transcription by modulating the expression of DNA methyltransferases (DNMTs), 5-methylcytosine(5mC) dioxygenases (TET) enzymes and through reactive oxygen species (ROS)-mediated pathways, thereby altering DNA methylation and microRNA (miRNA) expression^13 14 15 16^.

Given the long gestational window, the fetus can epigenetically adapt to environmental stressors, altering gene expression and organ development^17^. Impaired lung development in early life is one of the strongest predictors of poor lung function in later childhood and adulthood^18 19 20 21^. Our previous work demonstrated that early-life allergen exposure induces multigenerational epigenetic changes such as gene-specific DNA methylation in lung tissues that contribute to airway hyperresponsiveness (AHR)^22^. Identifying such epigenetic changes in accessible surrogate tissues, such as blood leukocytes, is also critical for translational asthma research, as they may reflect airway inflammation and serve as biomarkers of disease risk^23 24 25^. This study, conducted under the NIEHS-funded Toxicant Exposures and Responses by Genomic and Epigenomic Regulators of Transcription (TaRGET II) Consortium^26^, investigates the impact of early-life iAs exposure on lung health and its epigenomic consequences. By examining both target (lung) and surrogate (amniotic fluid, blood) tissues, we aim to uncover key epigenetic signatures linked to asthma risk. This work addresses a critical gap by investigating the legacy effects of toxic exposures on offspring health. It informs potential updates to WHO and EPA safety standards and supports environmental health equity. The public release of omics data from this study empowers the development of new epigenetic risk models across toxicology, developmental biology, and family health, especially in vulnerable populations. Moreover, these findings may guide the development of early intervention strategies to reduce asthma risk in affected communities, given the reversible nature of epigenetic modifications^27 28^.

## METHODS

### Animal Design

All animal experiments were approved by the Institutional Animal Care and Use Committees at Johns Hopkins University and University of Pittsburgh. Mice were maintained on AIN-93G diet (Research Diet, NJ) devoid of fishmeal or rice products (potential sources of arsenic) and given drinking water containing 0 or 10 ppb sodium arsenite (≥90% pure; Sigma-Aldrich) every other day. Both chow and water were batch-tested for arsenic content. Female nulliparous mice were exposed to 0 or 10 ppb iAs in water from two weeks before conception through lactation (eight weeks total), following TaRGET II guidelines. At weaning, total arsenic [iAs (III+V), MMAs (III+V), and DMAs (III+V)] in dam urine and litter plasma were measured via HG-CT-AAS and HG-CT-ICP-MS at Dr. Styblo’s lab (UNC). F1 mice were not exposed to iAs after weanling. Blood (surrogate) and lung (target) tissues were collected from male and female F1 mice at 3 weeks and 5 months for downstream analysis. A subset of 5-month-old F1 mice underwent allergen challenge to assess AHR.

### Allergen Challenge and Measurement of Allergen-Induced AHR

A subset of 5-month-old F1 mice were sensitized with 100 μg house dust mite (HDM, Greer Laboratories, Lenoir, NC) or saline (control) via intraperitoneal injection (i.p.) on Day 1. On Days 14, 17 and 21, HDM was administrated via intratracheal instillation (i.t.). HDM endotoxin was reduced using EndoTrap HD (Hyglos GmbH, Germany) according to the manufacturer’s protocol. On Day 23, mice were anesthetized, paralyzed, and ventilated using Flexivent (SCIREQ, Montreal, Canada). AHR was measured as the change in pulmonary resistance (cmH_2_O.s/mL) after methacholine (MCh; Sigma-Aldrich, St. Louis, MO) challenge. The protocol is illustrated in **Fig. 1A**.

**Fig. 1.**
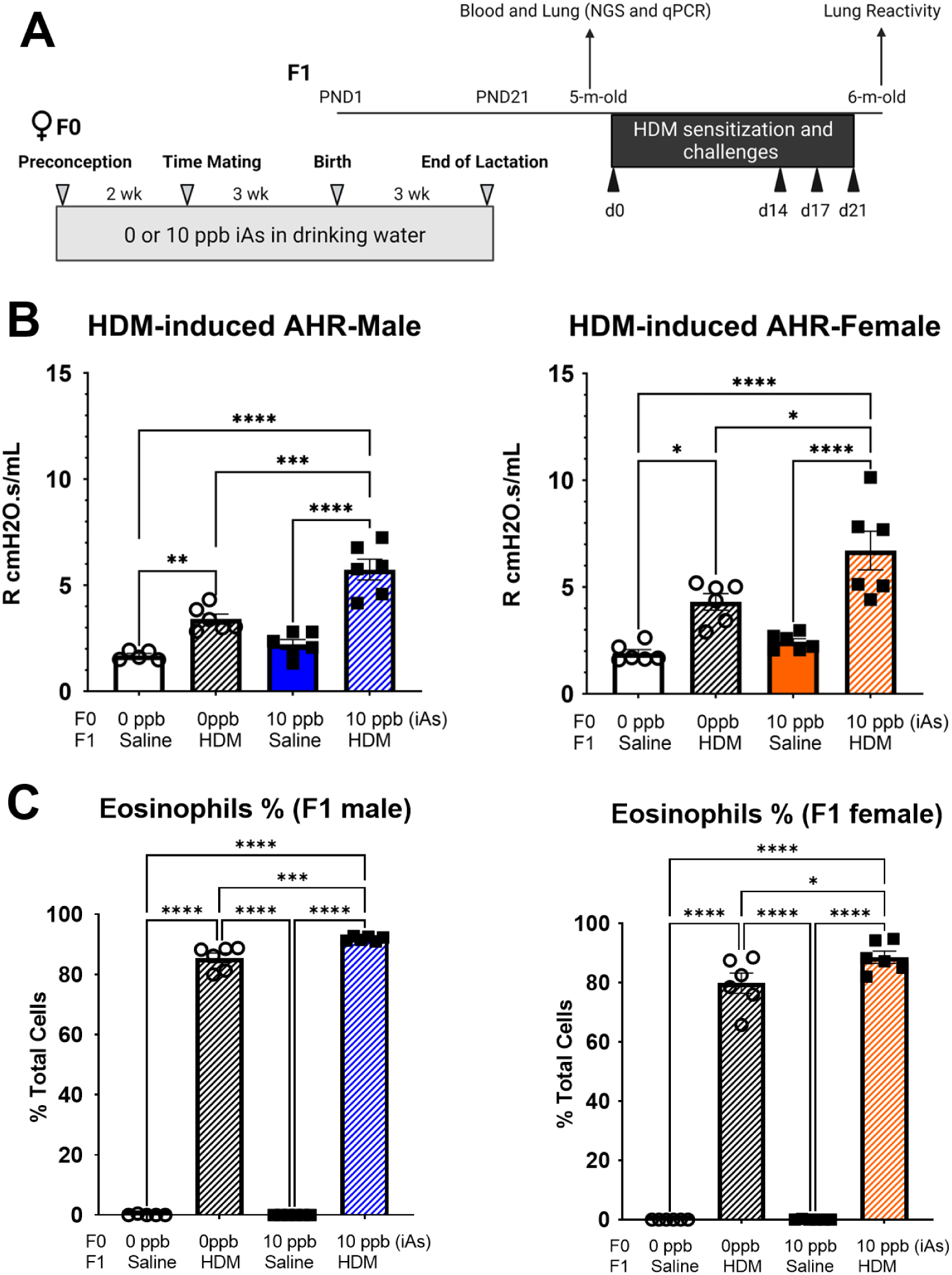
Maternal iAs exposure and study design to assess offspring asthma risk. A) Schematic shows the exposure window in which F0 dams received iAs in drinking water from preconception through lactation, followed by collection of F1 progeny for transcriptomic and methylomic analysis. A subset of 5-month-old F1 progenies were assessed for the HDM allergen-induced airway reactivity and airway inflammation. B) Lung resistance was assessed following methacholine (30mg/mL) challenge. C) BAL cells were stained with Diff-Quick, and eosinophil counts are shown. Each datapoint represents a pup from an individual dam; error bars indicate SEM. Five to six dams were used.

### Bronchoalveolar lavage (BAL) and Cell Counts

Following AHR measurements, BAL fluid was collected by flushing lungs with 1 mL saline containing cOmplete Mini Protease Inhibitor Cocktail (Roche, San Luis Obispo, CA). Total cell counts were performed immediately. Differential immune cell counts were assessed using cytospin slides stained with Diff-Quik.

### Tissue harvest, RNA and DNA isolation

Mice were euthanized at 3 weeks and 5 months of age with CO_2_. Blood and lung tissues were collected according to TaRGET II protocols. Lungs were snap-frozen and pulverized in liquid nitrogen for RNA and DNA extraction. Blood was collected via cardiac puncture into EDTA tubes and centrifuged to separate plasma and cells. Leukocytes were stored in Buffer RLT Plus (Qiagen) at −80°C. RNA and DNA from lung and blood were isolated using the AllPrep DNA/RNA/miRNA Universal Kit (Qiagen).

### Fetal Lung and Amniotic Fluid Collection

At embryonic day (E)16, fetal tissues were collected post-dam exsanguination using a modified Carraro et al^29^ protocol. Fetuses were dissected, and amniotic fluid was collected via insulin syringe after incising the amnion. Fetal lungs were flash-frozen and pulverized for nucleic acid extraction using the AllPrep Kit. miRNA from amniotic fluid was isolated using the NucleoSpin® miRNA Plasma Kit (Takara Bio, CA).

### RNA Sequencing and Transcriptomics Analysis

RNA-seq libraries from blood leukocytes of 5-month-old F1 mice were prepared per TaRGET II protocols. Libraries were sequenced at Genewiz using Illumina TruSeq Stranded Total RNA protocol on a HiSeq 10x platform. Paired-end reads were quality trimmed and mapped to the UCSC mm10 genome using CLC Genomics Workbench (Qiagen, Germantown, MD). Differential gene expression analysis was conducted using CLC’s DE feature; genes with fold change >2 and p < 0.05 were considered significant. Each sample yielded 25–50 million read pairs. Pathway analysis was performed with Ingenuity Pathway Analysis (IPA; Qiagen).

### Whole Genome Bisulfite Sequencing (WGBS) Analysis

Genomic DNA from blood leukocytes of 5-month-old F1 mice (DIN ≥ 7.0) was fragmented (550 bp insert size), end-repaired, and ligated to adapters before bisulfite conversion using the EZ-96 DNA Methylation Gold Kit (Zymo). WGBS libraries were generated using the Accel-NGS Methyl-Seq Kit (Swift Biosciences, Cat# 30096) and dual-indexed (Cat# 390384). Reads were quality-checked (fastqc^30^), trimmed (TrimGalore^31^) and aligned to the GRCm39 mouse reference genome using Bismark^32^. PCR duplicates were removed. Methylation calling was performed using the bismark_methylation_extractor with the following flags: -multicore 4, -cytosine_report, -ignore_r2 2, - ignore_3prime_r2 2, -bedGraph, -counts, -p, -no_overlap, and -report. Samples were deduplicated with deduplicate_bismark. Downstream analysis was done in R (v4.2.0) with Bioconductor packages. CpGs with ≥10× coverage, q ≤ 0.05, and ≥25% methylation difference were selected using *methylKit*. Differentially methylated CpGs were annotated using *methylKit*’s annotateFeatureFlank and mm39 BED/GTF files to assess CpG island/shore, gene region, and TSS proximity.

### Gene Expression Analysis

Total RNA (1 μg) isolated from lungs of E16, blood and lung tissues of 3-week-old, 5-month-old F1 mice was reverse transcribed with iScript Reverse Transcriptase (BIO-RAD, Hercules, CA). mRNA expression was quantified by SYBR Green-based qPCR, and relative expression was calculated by the 2^–ΔΔCt method, normalized to *rpl44* and universal mouse RNA reference (Qiagen). miRNA levels were quantified using the Mir-X miRNA qRT-PCR TB Green Kit (Takara Bio, CA). Primer sequences are listed in Supplementary Table 1 **(Supplementary Table 1)**.

### Plasma COL3A1 Protein Quantification

COL3A1 protein levels were measured in plasma from 3- and 5-month-old F1 mice using the Mouse Collagen III ELISA Kit (LS Bio, LS-F25411). Plasma was diluted 1:10 in Sample Diluent. Total protein was quantified using the Pierce BCA Protein Assay (Thermo Scientific), and COL3A1 values were normalized accordingly.

### Statistical Analysis

Relative gene expression and airway responsiveness data are presented as mean ± standard error of mean (SEM), with 5-6 mice per sex per treatment group (F1 males and females selected from individual dam). Technical triplicates were included for each assay. Statistical significance was assessed by ANOVA followed by Tukey’s multiple comparisons test. *P < 0.05, **P < 0.01, ***P < 0.001, ****P < 0.0001 vs. control. Data were analyzed and plotted using Prism 6 (GraphPad Software, La Jolla, CA, USA).

## RESULTS

### Impact of Maternal Exposure to iAs on Offspring’s Allergic AHR Risk

We investigated whether maternal exposure to the WHO guideline level of inorganic arsenic (iAs, 10 ppb) in drinking water influences offspring lung health. Female nulliparous mice were exposed to 0 or 10 ppb iAs in drinking water from two weeks before conception through lactation (eight weeks total), following TaRGET II Consortium guidelines. Importantly, F1 progeny were not exposed to iAs after weanling, allowing us to assess whether iAs effects on offspring lung health stemmed from maternal exposures. To evaluate offspring susceptibility, 5-month-old F1 mice were acutely challenged with HDM allergen **(Fig. 1A).** Both F1 males and females prenatally exposed to iAs exhibited significantly higher lung resistance compared to unexposed controls **(Fig. 1B).** Additionally, BALF from iAs-exposed offspring showed increased eosinophil proportions **(Fig. 1C)**. These findings indicate that the maternal iAs exposure increases offspring susceptibility to allergen-induced airway inflammation and reactivity. While iAs exposure alone did not significantly induce allergic responses or impair lung function, it primed offspring for enhanced AHR upon later-life HDM challenges.

### Long-Lasting Changes in Blood Transcriptome and Methylome after Early-Life iAs Exposure

To investigate the molecular mechanisms underlying this increased susceptibility, we profiled gene expression in blood leukocytes from 5-month-old F1 mice prior to HDM challenge. RNA-seq revealed 642 differentially expressed genes (DEGs) in males and 599 in females, with 70 overlapping **(Fig. 2A and Supplementary Table 2)**. Ingenuity pathway analysis (IPA) revealed that overlapping DEGs were enriched in acute phase response and immune pathways (e.g., IL-1, IL-12), while male-specific DEGs included LXR/RXR and NRF2 signaling, and female-specific DEGs involved pulmonary fibrosis and airway pathology pathways. Analysis of upstream regulators and networks identified TGFβ as the top regulator linking DEGs (common in F1 male and female) involved in extracellular matrix (ECM) remodeling (e.g., *arg1, bmp1, col3a1, itih3 map1b, tnc* and *tpsb2*), inflammation (e.g., *fga, fgb, f2, hamp, mlxipl* and *saa3*), and embryonic development (e.g., *afm, alb, aldob, igfbp5*, and *serpina1b*) **(Fig. 2B)**. These findings suggest that maternal iAs exposure primes immune and tissue development pathways in blood leukocytes, through modifications in leukocyte transcriptome, possibly influencing later respiratory outcomes.

**Fig. 2.**
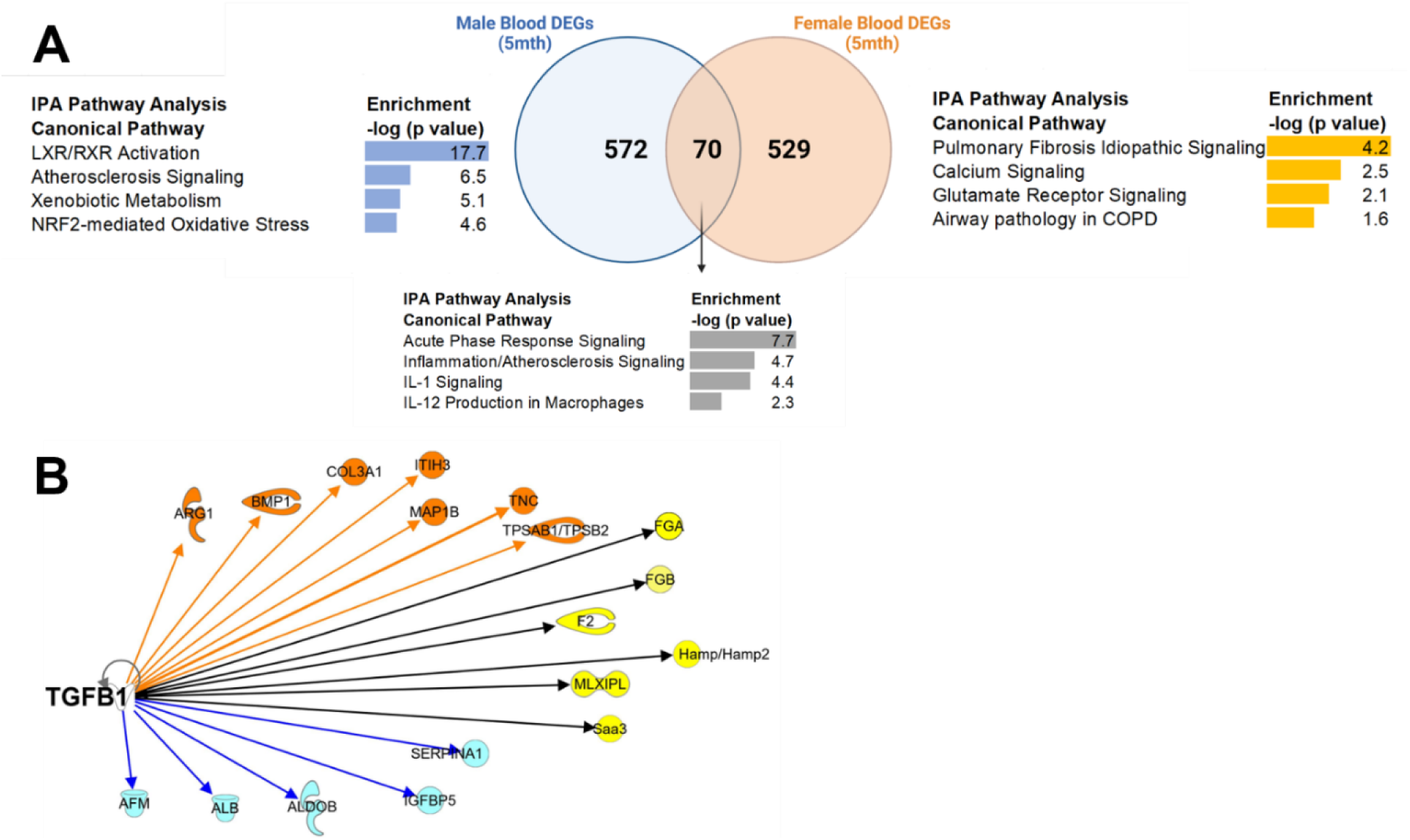
Blood transcriptome profiles and predicted canonical pathways and regulatory networks. A) Numbers of differentially expressed genes (DEGs) identified in male and female progeny are illustrated with a Venn diagram. Enriched canonical pathways ranked by the p value are shown. B) Using the set of DEGs common in both sexes, Ingenuity Pathway Analysis (IPA) highlights TGFβ as a central upstream regulator linking overlapping DEGs related to extracellular matrix signaling (orange), inflammation (yellow), and embryonic development/cell metabolism (blue).

To determine whether iAs induces the long-lasting DNA methylation changes that modulate blood transcriptome, we conducted WGBS on the same blood leukocytes we profiled the transcriptome. We identified 24 differentially methylated CpGs (DMCs) in males and 17 DMCs in females **(Fig. 3A and Supplementary Table 3),** with four DMCs overlapping annotated to *miR-101c*. *miR-101c* exhibited hypermethylation **(Supplementary Fig. 3A)** and was significantly downregulated (∼6-fold in males, ∼2-fold in females; **Fig. 3B**). This suggests that iAs-induced DNA methylation changes at *miR-101c* are associated with its downregulation. To further explore the functional significance of *miR-101c,* we conducted pathway analysis to examine its relationship with TGFβ-connected DEGs **(Fig. 3C)**. Although none of TGFβ-connected DEGs were direct targets of *miR-101c*, some (i.e., *mlxipl, tnc, serpina1b*, *col3a1, bmp1, map1b, itih3, alb* and *fga*) may be indirectly regulated by *miR-101c*. These findings suggest *miR-101c* downregulation contributes to iAs-induced transcriptional reprogramming.

**Fig. 3.**
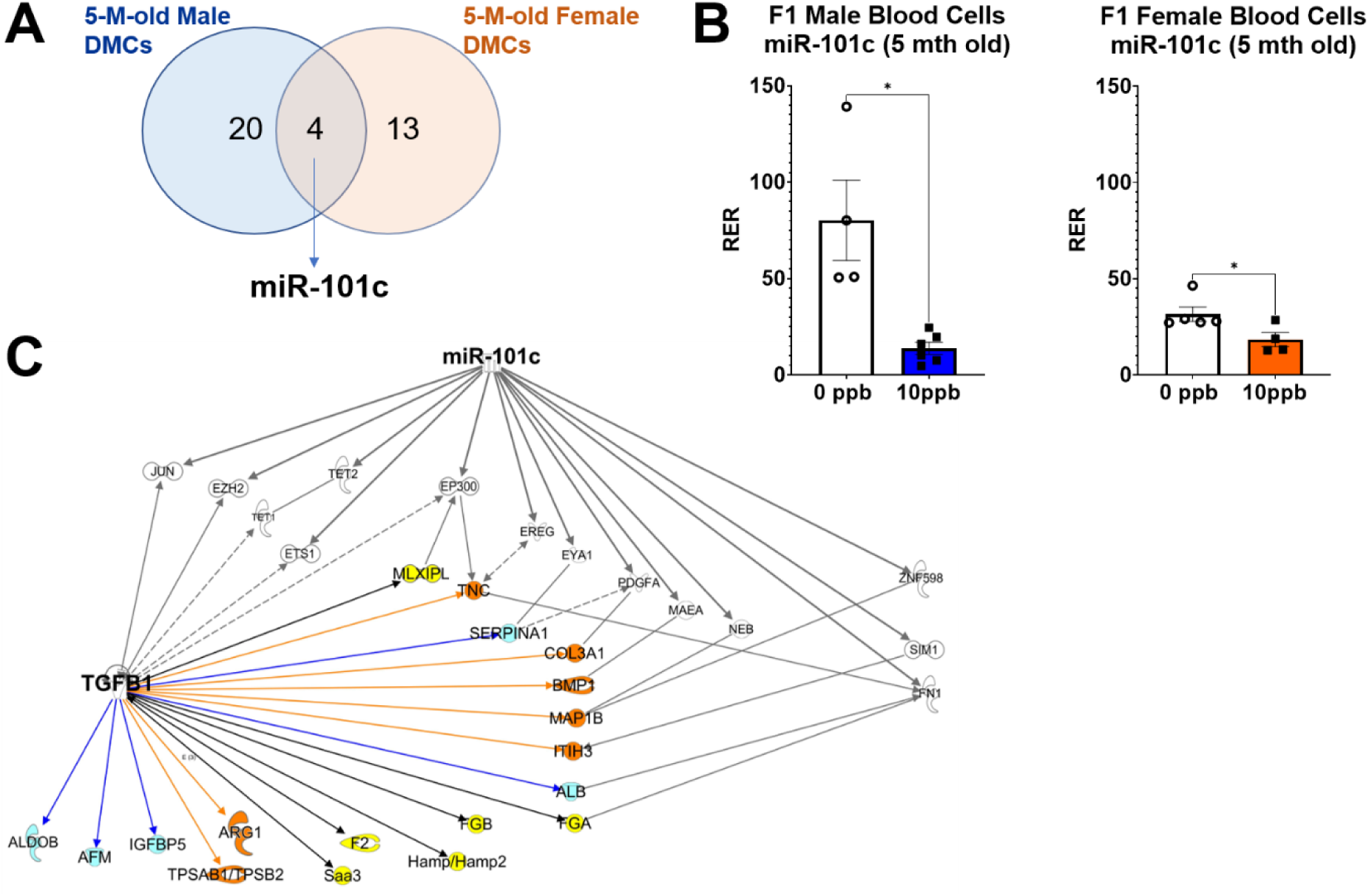
Blood methylome analysis and associations between *miR-101c* and TGFβ-related differentially expressed genes. A) Number of differentially methylated CpGs (DMCs) identified in male and female progeny are illustrated with a Venn diagram. There are four DMCs overlapping DMCs annotated to *miR-101c*. B) Expression levels of *miR-101c* are shown. C) IPA-derived network linking this miRNA to TGFβ-related DEGs. Each datapoint represents a pup from an individual dam; error bars indicate SEM. Four to five dams were used.

To assess developmental persistence, we compared blood transcriptomes of 3-week-old (juvenile) and 5-month-old (adult) progeny **(Supplementary Table 4).** *miR-101c* was consistently downregulated across age and sex. Several TGFβ-related DEGs also persisted from juvenile to adult blood leukocytes, including *col3a1* and *fga* in both sexes, while others (e.g., *mlxipl, saa3*) showed opposite expression trends across ages. These dynamic shifts suggest that iAs-induced epigenetic changes may be modulated by developmental stage or sex-specific physiology.

### Blood Transcriptome as Predictor of Later-life AHR

We next evaluated whether early blood gene expression predicted later AHR phenotypes. Correlations between blood DEGs (pre-HDM) and AHR (post-HDM) were assessed **(Fig. 4A and Supplementary Table 5)**. In males, early expression of *col3a* (r=0.88, p=0.02) and *fgb* (r=0.83, p=0.04) were positively associated with AHR, while *miR-101c* (r=-0.79, p=0.06) showed a negative correlation. These associations persisted into adulthood (*col3a* r=0.92, *fgb*, r=0.83 and *miR-101c*, r=-0.81). In females, *col3a1* expression at both 3 weeks (r=0.91) and 5 months (r=0.92) positively correlated with AHR. *miR-101c* expression remained inversely correlated with AHR risk across the ages (3 weeks, r=-0.87 and 5 months, r=-0.84). Some correlations (e.g., *f2* at 3 weeks, *map1b* and *igfbp5* at 5 months) were age-dependent, particularly in females, suggesting developmental modulation of gene-AHR relationships.

**Fig. 4.**
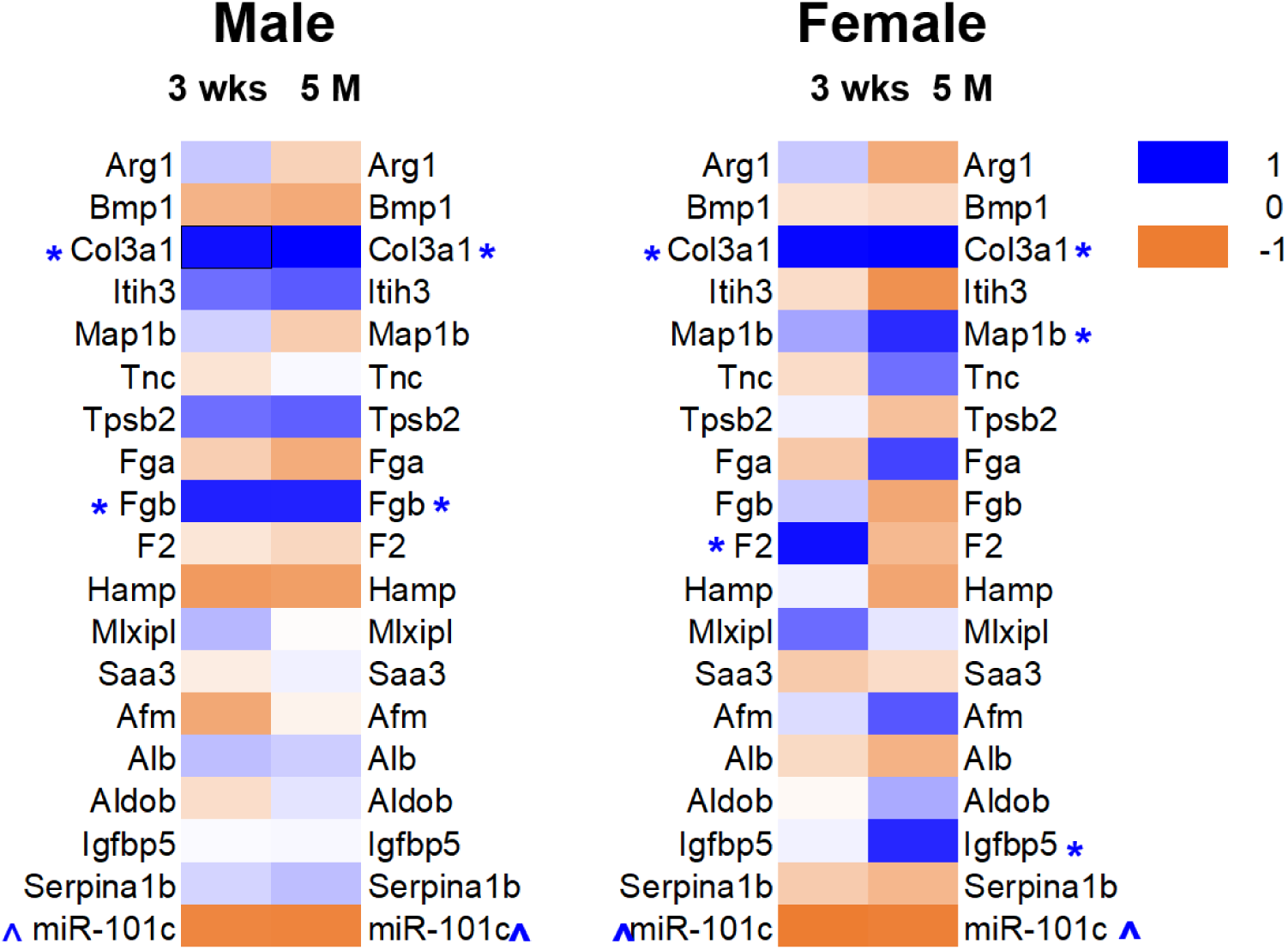
Relationship between iAs-induced blood transcriptional changes and allergen-induced AHR. This figure presents correlation patterns between baseline blood DEGs expression changes prior to HDM challenge and subsequent changes in AHR. Pearson correlation coefficients are depicted using a color gradient to distinguish positive (blue) and negative (peach) associations. * Positive correlation: r>0.8, p<0.05; ^ negative correlation: r<-0.8, p<0.05

To validate these molecular markers for maternal iAs exposure, we measured plasma levels of *miR-101c* and COL3A1 protein. Plasma *miR-101c* was reduced in iAs-exposed F1 mice (both sexes) across ages **(Fig. 5A)**, while COL3A1 protein increased in 5-month-old plasma and showed an upward trend in juveniles **(Fig. 5B).** These findings support their use as biomarkers of maternal iAs exposure and asthma risk if validated in human population cohorts of iAs-exposed community.

**Fig. 5.**
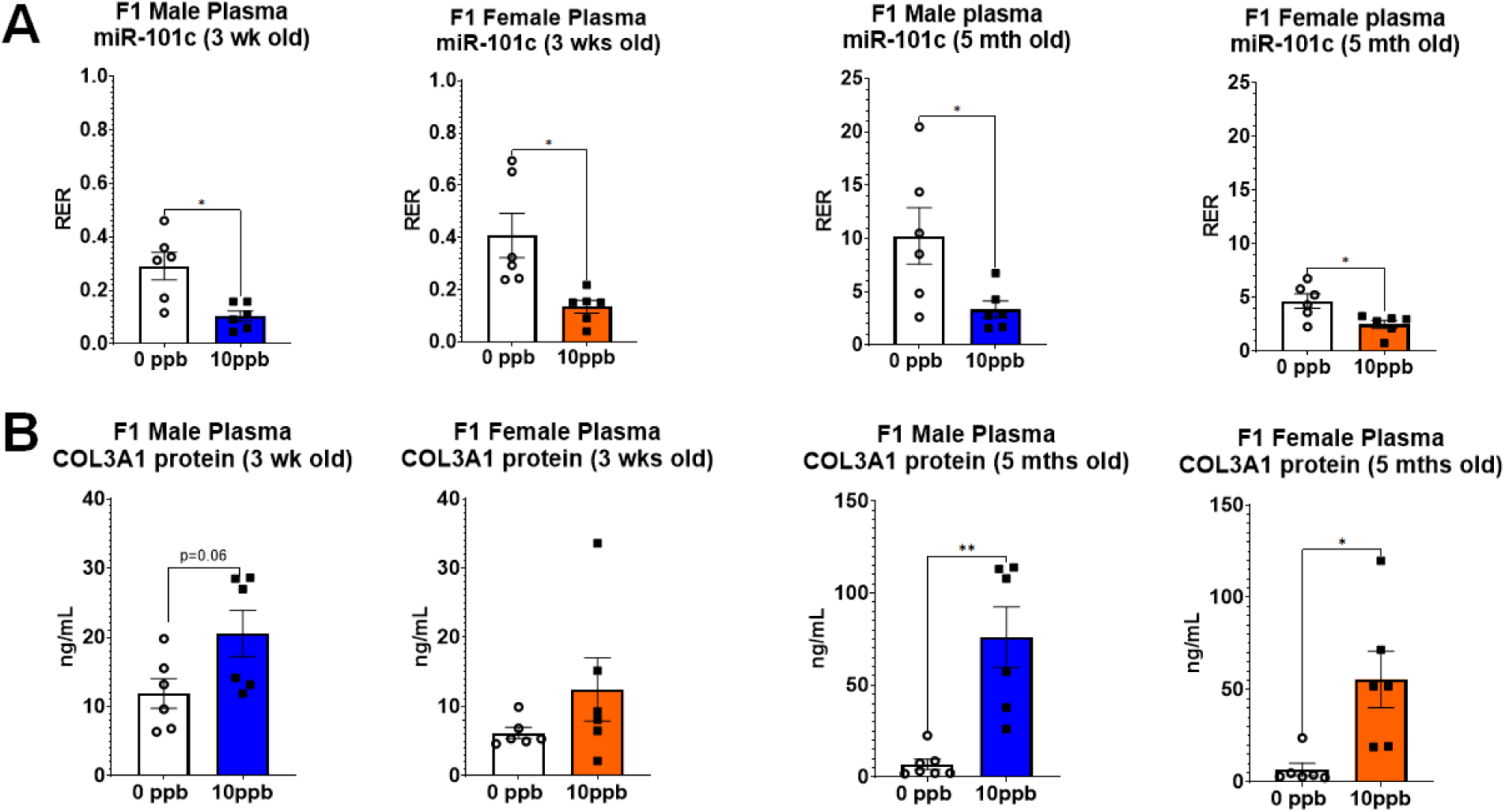
Circulating miR-101c levels in progeny exposed to maternal iAs. Plasma levels of miR-101c and COL3A1 protein are shown at different developmental stages. A) Relative expression of miR-101c across F1 progeny at juvenile (3 weeks of age) and adult (5 months of age) prenatally exposed to iAs. B) Levels of COL3A1 protein are shown at juvenile and adult F1 progeny. Each datapoint represents a pup from an individual dam, error bars indicate SEM.

### Overlap Between Blood and Lung Transcriptional Changes is Time- and Sex-Dependent

We next examined whether iAs-associated transcriptional changes observed in blood were also present in lung tissues. In juvenile males, 5 out of 18 TGFβ-mediated DEGs (solid black bar, p <0.05) were differentially expressed in lung tissue **(Fig. 6A and Supplemental Table 4)**, with *fga* and *fgb* upregulated in both blood and lung. However, some DEGs (e.g., *itih3*, *tpsb2)* showed opposite directions between tissues. *Col3a1* was upregulated only in lung tissue. In adult males, 9 TGFβ-related DEGs (solid black bar, p < 0.05) were altered in lungs **(Fig. 6B and Supplemental Table 4).** Seven DEGs (*col3a1, map1b, tpsb2, f2, saa3, aldob*, and *serpina1b*) were upregulated in both blood and lung, while *itih3* and *hamp* were upregulated in blood but downregulated in lung. Strikingly, no DEGs showed consistent changes across tissue and developmental stages, though *fga* and *fgb* were consistently upregulated in both tissues with near statistical significance.

**Fig. 6.**
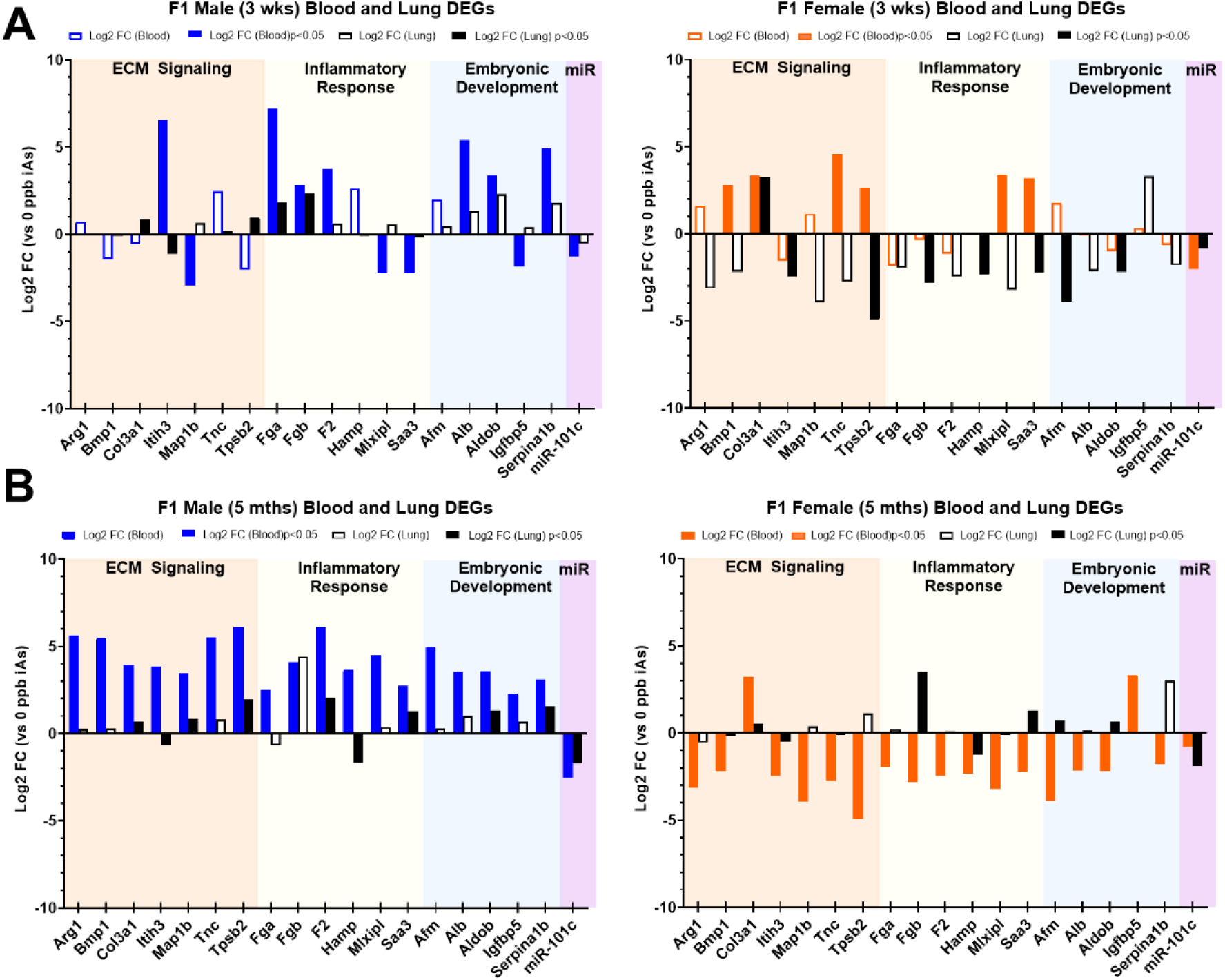
Comparison of transcriptional patterns across tissues and developmental stages. Expression changes (FC: fold changes vs. unexposed) of DEGs and *miR-101c* are compared between blood and lung tissues in both A) juvenile (3 weeks of age) and B) adult (5 months of age) progeny. Blue and orange bars represent FC in blood from male and female progeny, respectively. Black bars represent FC in lung tissues. Solid bars indicate statistically significant changes.

In juvenile females, 8 TGFβ-related genes (solid black bar, p < 0.05) were differentially expressed in lungs **(Fig. 6A and Supplemental Table 4)**. Only 3 DEGs (*col3a1, tpsb2*, and *saa3*) matched the direction in blood. Others (e.g., *itih3*, *hamp, fgb*, *afm*, and *aldob*) showed discordant patterns. In adult females, 8 TGFβ-related genes (solid black bar, p < 0.05) were altered in lung tissues (**Fig. 6B and Supplemental Table 4)**, with *col3a1* upregulated and *itih3* and *mlxipl* downregulated in both blood and lung. Other DEGs (e.g., *tpsb2, fgb, saa3, afm,* and *aldob*) in lung showed opposite direction in blood. Notably, *col3a1* was the only DEG consistently upregulated in lung tissues across developmental stages in both sexes.

*miR-101c* downregulation was also observed in lung tissues of iAs-exposed mice. Both juvenile and adult lung tissues from males and females showed decreased *miR-101c* expression, as similarly seen in blood **(Fig. 6A and 6B)**. These findings highlight *miR-101c* as a stable epigenetic marker of maternal iAs exposure in both target (lung) and surrogate (blood) tissues regardless of sex.

### Epigenetic Programming of miR-101c Occurs Early in Gestation

Given the persistent downregulation of *miR-101c*, we assessed its expression in embryonic lungs at E16. *miR-101c* levels were significantly reduced in lung buds of female fetuses prenatally exposed to iAs and showed a downward trend in males **(Fig. 7A)**. We also examined *miR-101c* levels in amniotic fluid (AF), which plays a key role in fetal lung development. In unexposed mice, males had lower AF *miR-101c* levels than females. Upon iAs exposure, *miR-101c* levels decreased in AF from both sexes **(Fig. 7B).** Although the link between *miR-101c*, its regulatory effects on target genes, and fetal lung development requires further validation, our findings suggest that *miR-101c* epigenetic programming begins early in gestation and persists in postnatal life, potentially mediating long-term susceptibility to respiratory disease.

**Fig. 7.**
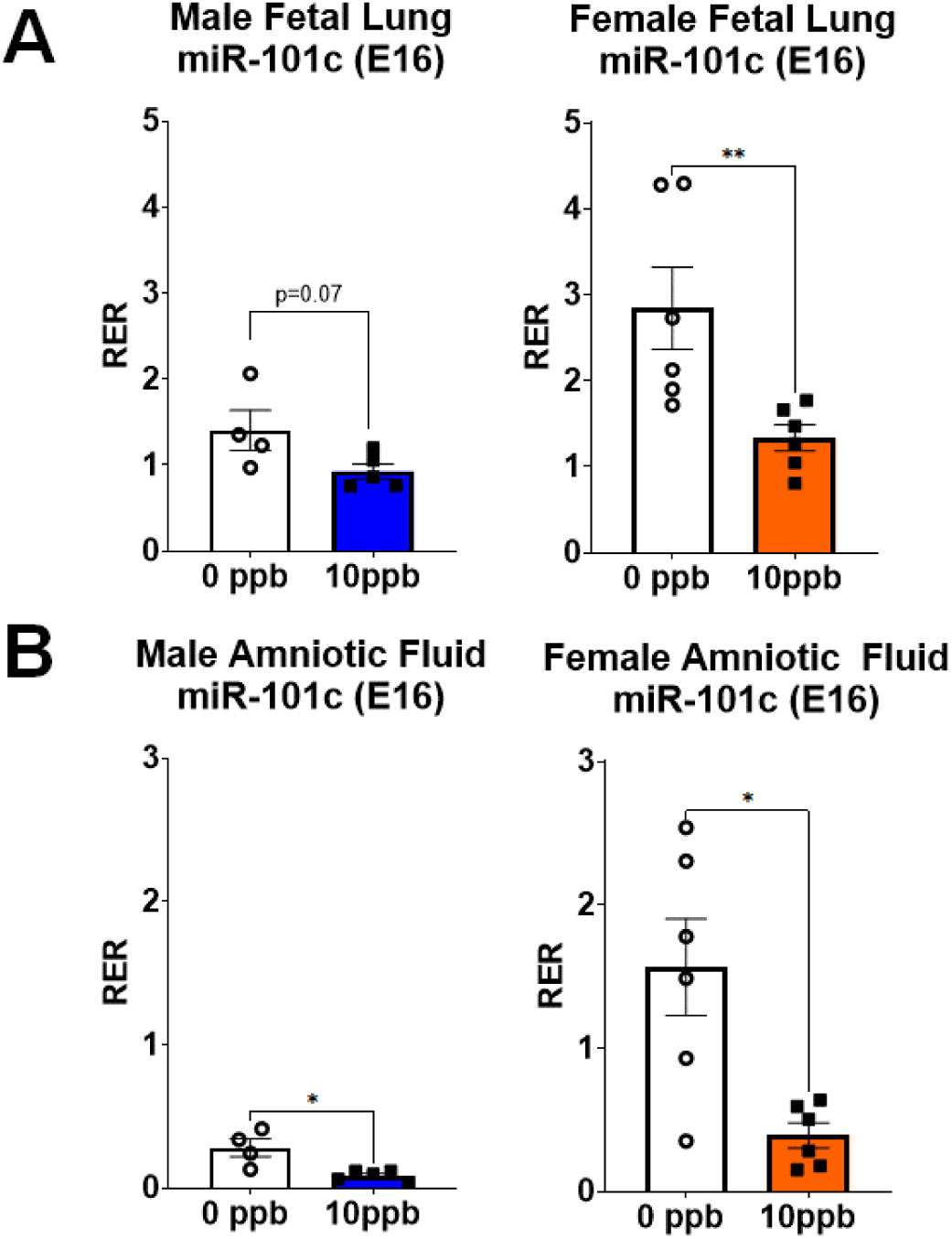
Fetal lung and amniotic fluid levels of miR-101c. Relative levels of *miR-101c* are shown in fetal lung tissues and matched amniotic fluid at embryonic day 16. Data illustrate expression patterns in the context of maternal iAs exposure. Each datapoint represents a pup from an individual dam, error bars indicate SEM.

## DISCUSSION

In this study, we demonstrated the epigenetic impact of early-life exposure to 10 ppb iAs (the current WHO and EPA provisional limit in drinking water) on offspring’s long-term lung health using an experimental mouse model of asthma. Our previous work showed that *in utero* allergen exposure leads to gene-specific DNA methylation changes, promoting more severe allergic disease^22^, suggesting early-life epigenetic reprogramming from exposures determines offspring’s asthma risk. Here, we investigated the adverse effects of low-dose iAs exposure during early development on asthma risk. Unlike previous studies using higher iAs doses (>100 ppb), which have demonstrated roles in cancer and other diseases like cardiovascular and neurological disease, our study focused on 10 ppb iAs, delivered only *in utero*. We found no significant effect of this exposure on baseline lung function. However, upon acute HDM challenges in adulthood, both male and female offspring from iAs-exposed mothers exhibited greater lung resistance compared to unexposed controls. This supports the “second hit hypothesis” suggesting that maternal iAs exposure may prime the offspring to be more vulnerable to allergic airway inflammation later in life.

In humans, iAs has been detected in cord blood at delivery, suggesting transplacental transfer, accumulation in fetal tissues, and potential impacts on fetal development. However, there are mixed findings regarding the association between iAs level and fetal growth^33 34 35 36^. Some studies indicated that maternal arsenic metabolism increases during pregnancy, potentially protecting the fetus by reducing iAs and MMA exposure^37 38^. Additionally, arsenic level in breast milk remains low even when maternal drinking water is highly contaminated with As^39 40^. In our mouse model, F1 progeny exposed *in utero* to iAs showed a 20-fold lower arsenic level (0.22 ppb) compared to dams (4.6 ppb) **(Supplementary Fig 1)**, indicating minimal transfer. Moreover, no significant differences in birth weight or postnatal growth were observed **(Supplementary Fig. 2)**, suggesting that asthma risk was not attributable to generalized developmental toxicity of low dose *in utero* iAs exposure.

Indeed, we found that early-life exposure to iAs exerts a lasting effect on gene transcription in both blood and lung tissues of adult (5-month-old) mice. Of note, F1 progeny received no further iAs exposure after weanling, indicating that the transcriptomic changes established from prenatal exposures. Our findings suggest that early-life exposure to iAs induces epigenetic regulation of gene transcription, priming the offspring’s lung to be more susceptible to AHR upon later allergen challenges. This hypothesis is supported by RNA-seq analysis of blood leukocytes from adult F1 progeny, which revealed significant transcriptional changes in genes linked to inflammatory response. Although beyond the scope of this study, these long-lasting transcriptomic effects may also contribute to other inflammatory diseases. When comparing transcriptional changes in TGFβ-related DEGs in both sexes at 3 weeks (just after maternal iAs exposure) and 5 months (after 4 months without iAs exposure), not all these DEGs showed consistent expression patterns over the course of development or correlations with allergen-induced AHR. These findings indicate that other developmental factors, such as hormonal or environmental influences, may modify iAs-induced transcriptional programs over time. Further studies are required to identify these modulators.

We next examined mechanisms underlying these persistent transcriptional changes. Gestation and early-life are periods of enhanced epigenetic vulnerability, characterized by genome-wide DNA demethylation, remethylation, and the establishment of tissue-specific methylation patterns^41^. MiRNAs^42^ are also hypothesized to modulate the maintenance of existing epigenetic signatures and the establishment of new patterns of DNA and histone modifications in somatic and germ cells by targeting epigenetic regulators^43 44^. Using WGBS, we identified *miR-101c* as the only consistently differentially methylated in blood leukocytes from both male and female progeny of iAs-exposed mothers. This methylation change corresponded with reduced *miR-101c* expression. Therefore, the downregulation of *miR-101c* may influence the expression and/or activity of its downstream targets. Because no DNA methylation changes were observed in the DEGs identified by RNA-seq, we postulate that persistent transcriptional reprogramming may be mediated by other epigenetic mechanisms or indirectly via *miR-101c* regulation, as indicated by potential connected pathways. Strikingly, downregulation of *miR-101c* was detected during late gestation (in E16 lung buds and amniotic fluid) and persisted postnatally in both blood and lung tissues at 3 weeks and 5 months. The regulation *of miR-101c* in lung development and its sensitivity to iAs exposure warrants further exploration.

The transcriptomic and methylomic data analyses provide new insights into respiratory health in populations exposed to contaminated drinking water and highlight potential biomarkers for iAs risk assessment. Pathway analysis of blood RNA-seq data identified TGFβ signaling as a key upstream network connecting DEGs involved in extracellular matrix (ECM) remodeling, embryonic development, and inflammatory responses. Dysregulation of these pathways could increase allergen-induced AHR by modulating the structural matrix and inflammatory state of the airway. Correlation analysis further linked transcriptional changes in DEGs such as *col3a1*, *fgb*, *map1b*, *igfbp5* and *miR-101c* with AHR risk, although the direction and degree of these associations varied by sex. Importantly, these transcriptional changes persisted even after the cessation of iAs exposure, suggesting stable epigenetic programming.

Using the Public Health Genomics and Precision Health Knowledge Base v10, we explored the links between selected DEGs, human diseases and environmental exposures. *FGB* encodes the β-chain of fibrinogen. *FGB* genetic variants are associated with metabolic disorders^45^ and myocardial infarction^46 47^, and can modulate IL-6 responses to air pollution in cardiovascular disorders^48 49^. *COL3A1* encodes collagen type III protein, a major ECM component in skin, lung and blood vessels. Variants in *COL3A1* have been associated with cardiovascular diseases^50^ and serve as prognostic markers in melanoma^51^. A review of public databases indicates that *MiR-101-3p* (the human homolog of mouse *miR-101c*) is linked to cognitive impairment^52^ and other stress-related outcomes. Our results show that blood *col3a1* and *miR-101c* levels strongly correlate with AHR risk. Maternal iAs exposure increased plasma COL3A1 protein levels and decreased *miR-101c* expression of both plasma and amniotic fluid. These findings suggest *col3a1* and *miR-101c* could serve as molecular signatures of prenatal iAs exposure in surrogate tissues (e.g., blood or amniotic fluid), potentially informing strategies for risk assessment of iAs exposure if validated in independent human cohorts.

We next validated overlapping DEGs in paired lung tissues to test whether early-life iAs exposure establishes similar epigenetic programs in both blood and lung. Aberrant epigenetic signatures in lung tissues can disrupt genes critical for asthma pathogenesis. We observed consistent upregulation of *col3a1* and downregulation of *miR-101c* in both blood and lung tissues across juvenile and adult progeny of iAs-exposed mothers, regardless of sex. COL3A1 plays a central role in ECM remodeling and airway fibrosis, processes closely linked to asthma severity^53^. A previous study reported that *in utero* exposure to moderate iAs levels (50 and 100 ppb) increased AHR and *col3a1* expression in mouse offspring. Remarkably, these effects persisted even when postnatal exposure ceased, supporting the concept that early-life iAs exposures establish a lasting imprint on lung function^54^. In other study, overexpression of COL3A1 was observed in lung fibrosis and its aberrant expression was regulated by histone deacetylation^55^, suggesting *COL3A1* is epigenetic regulated. In this study, we found that maternal exposure to low dose of iAs is associated with decreased DNA methylation at the *col3a1* promoter **(Supplementary Fig. 3B)**. Collectively, targeting COL3A1 through epigenetic modulators such as histone deacetylase inhibitor (HDACi), may help counteract iAs-induced airway remodeling.

*MiR-101*^56^ is recognized as a tumor suppressor and may promote osteogenic differentiation of bone marrow mesenchymal stem cells by targeting EZH2/WNT signaling pathway^57^. Its role in asthma, however, remains understudied. One report showed that plasma *hsa-miR-101* was downregulated in asthmatic patients after rhinovirus stimulation, correlating with interferon signaling^58^, suggesting a role for *miR-101* in regulating lung immune responses to viral infection. Additionally, *miR-101* has been implicated in the NFkβ pathway, which can be modulated by heavy metal exposures^59^. To understand its relevance in lung health, we explored genes potentially targeted by *miR-101*. Beyond EZH2, *miR-101* may regulate epigenetic enzymes such as TET1 and TET2, and transcription factors like EP300, JUN, and ETS-1. Integrative analyses of iAs-affected DEGs and *miR-101c* suggest potential regulatory links. We speculate that downregulation of *miR-101c* could increase PDGFA expression, subsequently enhancing COL3A1 expression and tissue remodeling^60^, though this requires further validation. Furthermore, we found maternal iAs exposure was associated with downregulation of *miR-101c* in fetal lung buds and amniotic fluid, suggesting alterations in lung or circulating *miR-101c* during gestation may affect organ development. Notably, this downregulation persisted in lung tissues through juvenile and adult stages, supporting a role for *miR-101c* in regulating genes involved in TGFβ and WNT signaling, immune function, and epigenetic modification.

Our findings also underscore the importance of incorporating sex as a biological variable in omics studies^61^. Transcriptomic and methylomic responses to early-life iAs exposure diverged by sex. The number of DEGs common to male and female adult blood (5-month-old) was limited (70), compared to sex-specific DEGs (572 in males, 529 in females). Similarly, *miR-101c* was the only one gene differentially methylated in both sexes. After accounting for developmental changes between 3 weeks and 5 months, only two genes (*mlxipl* and *saa3*) were differentially expressed in both sexes, interestingly, in opposite directions. In the lungs, only *col3a* and *tpsb2* were differentially expressed in both sexes at both juvenile and adult stages. Overall, *miR-101c* was the only gene consistently and differentially expressed across tissues, sexes and time points.

Although sex differences in asthma are well documented^62^, especially across the lifespan, this study focused on generalizable impacts of early-life exposure. Both male and female offspring from iAs-exposed mothers exhibited increased HDM-induced AHR in adulthood. Still, the underlying mechanisms may differ by sex. While both sexes showed similar AHR and inflammatory responses, they may result from distinct regulatory pathways. Sex hormones, alone or interacting with environmental factors, can influence gene transcription and epigenetic regulation^63 64 65^, potentially resulting in sex-specific trajectories toward a shared disease phenotype^66^. For example, in females, puberty-associated hormonal changes may shift gene expression within the same tissue. Except *Col3a*, most TGFβ-related DEGs in female lung tissue showed distinct expression profiles between juvenile and adult stages. Moreover, in females, we observed inconsistencies in the magnitude and direction of gene expression changes between blood and lung tissue, potentially explaining the differential correlation between DEGs and AHR risk across developmental stages. These discrepancies were less apparent in males, where transcriptional changes, especially in TGFβ-related genes, appeared more stable after the cessation of maternal exposure. We speculate that the absence of ovarian hormone fluctuations may contribute to this difference. Our omics analyses provide a foundation for interpreting arsenic-related epigenomic and epidemiologic data, though future research is needed to explore sex-specific regulatory mechanisms.

This study has limitations that warrant future investigation. While our data suggest that 10 ppb iAs, a concentration currently permitted under WHO and EPA standards, can exert long-term health impacts, we recognize the need to determine the lowest iAs exposure level without adverse effects. In the future, our findings should be compared to published epigenomic and epidemiological data to help establish predictive models assessing health risks at varying iAs concentrations. We analyzed transcriptome and methylome profiles at 3 weeks and 5 months of age. Future work should validate these changes across additional life stages including fetal development, early postnatal periods, and older age, given the dynamic nature of the epigenome throughout life^67^. MiRNAs^68^, including *miR-101c*, may disrupt existing epigenetic marks. Given their role as both biomarkers and mediators in childhood asthma^69^, incorporating miRNA profiling into longitudinal human studies could enhance understanding of disease development. Our study does not resolve cell-type-specific epigenetic effects. Single-cell transcriptomic and methylomic analyses could help identify transcriptional changes in specific lung cell populations (e.g., epithelial or immune cells), enabling a clearer connection between molecular alterations and functional outcomes. These high-resolution datasets may benefit from computational tools and machine learning approaches to model interactions between time- and cell-specific epigenetic programs and AHR phenotypes. Moreover, our current design does not pinpoint the critical window of vulnerability for iAs-induced epigenetic changes. It remains unknown whether life-long exposure to iAs magnifies transcriptional changes and AHR risk, with or without secondary insults. Exploring varied exposure windows, epigenomic profiles, and long-term health outcomes, including multi-generational inheritance, represents a crucial future direction, particularly in cohorts with longitudinal follow-up data. Whether epigenetic changes induced by iAs in blood and lung tissues are heritable remains unexplored.

In summary, this study provides novel insights into how early-life exposure to iAs reprograms the epigenome and increases susceptibility to asthma. *miR-101c* and COL3A1 emerged as persistent molecular signatures in blood, lung, and amniotic tissues, suggesting their potential utility as biomarkers for early identification of at-risk individuals. These findings could inform the development of targeted interventions and support policy efforts to ensure safe drinking water^70^ and reduce iAs exposures, especially for vulnerable communities.

## ACKNOWLEDGEMENTS

This work was supported by NIEHS as part of Toxicant Exposures and Responses by Genomic and Epigenomic Regulators of Transcription II (TaRGET II) Consortium through U01ES026721. We acknowledge program leadership by members of the NIEHS TaRGET II workgroups, especially Fred. L. Tyson, Kim McAllister, Christopher G. Duncan, Amanda Garton, Lisa H. Chadwick and Maya Evanitsky.

## AUTHOR CONTRIBUTIONS

Y.S., Q-y. S., H-y. Y., and J.P. contributed experiment execution and data analysis; M.W., D.R., B.S., and B.P. contributed multi-omic analysis; A.B., W.M., S.B., and W-y.T. contributed experimental design and result interpretation; and Y.S., D.R., and W-y.T contributed manuscript writing and A.B., W.M., S.B., and W-y.T contributed manuscript review.

---

**Supplementary Fig. 1.**
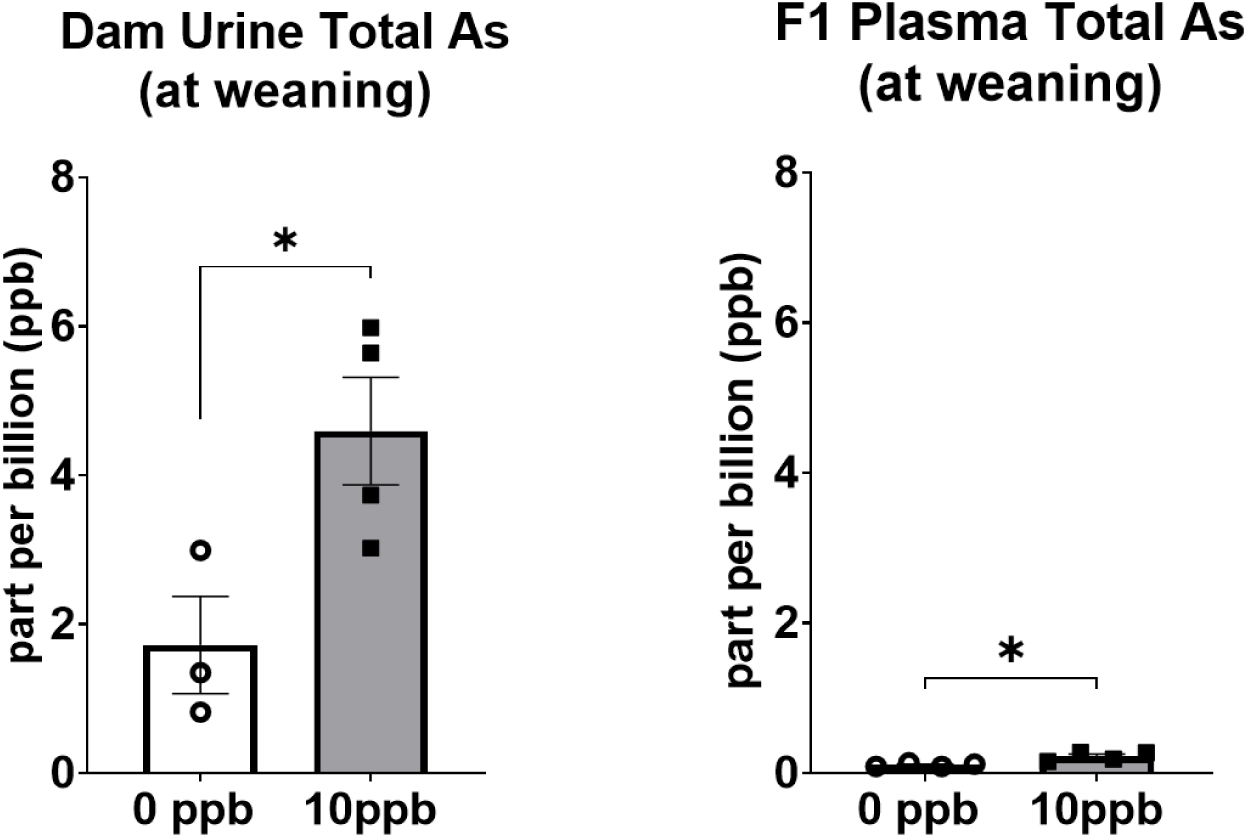
Circulating iAs metabolites in dam and pup. **T**otal As metabolites including iAs (III+V), MMAs (III+V) and DMAs (III+V) were assessed in maternal urine and plasma F1 progeny at weaning. Each datapoint represents an individual dam or a pup from an individual dam, error bars indicate SEM.

**Supplementary Fig. 2.**
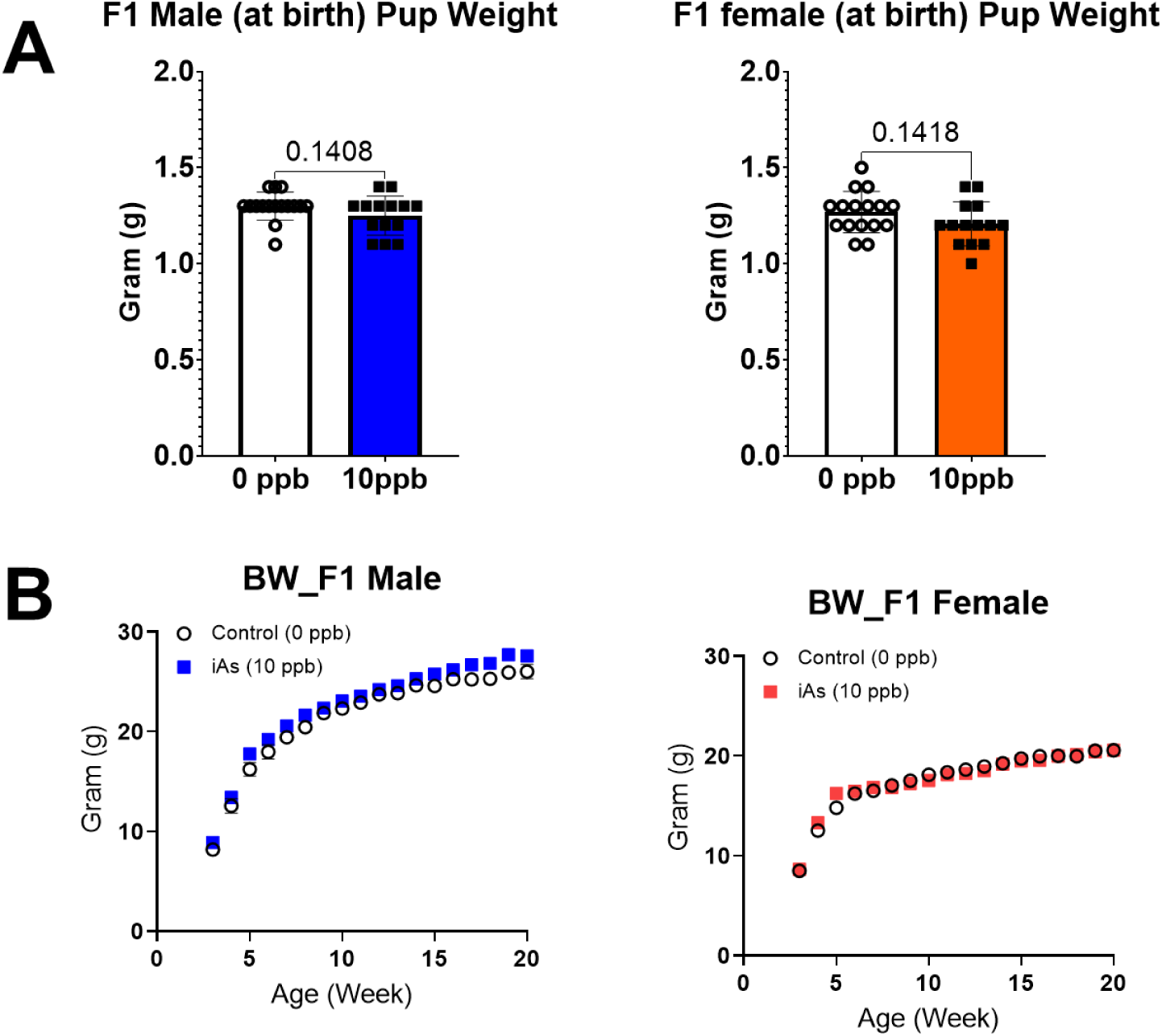
Body weight of F1 progenies across development. A) Each datapoint represents the birth weight of a pup from an individual dam at birth. B) Average body weight of 6-8 pups from all dams is shown. Error bars indicate SEM.

**Supplementary Fig. 3.**
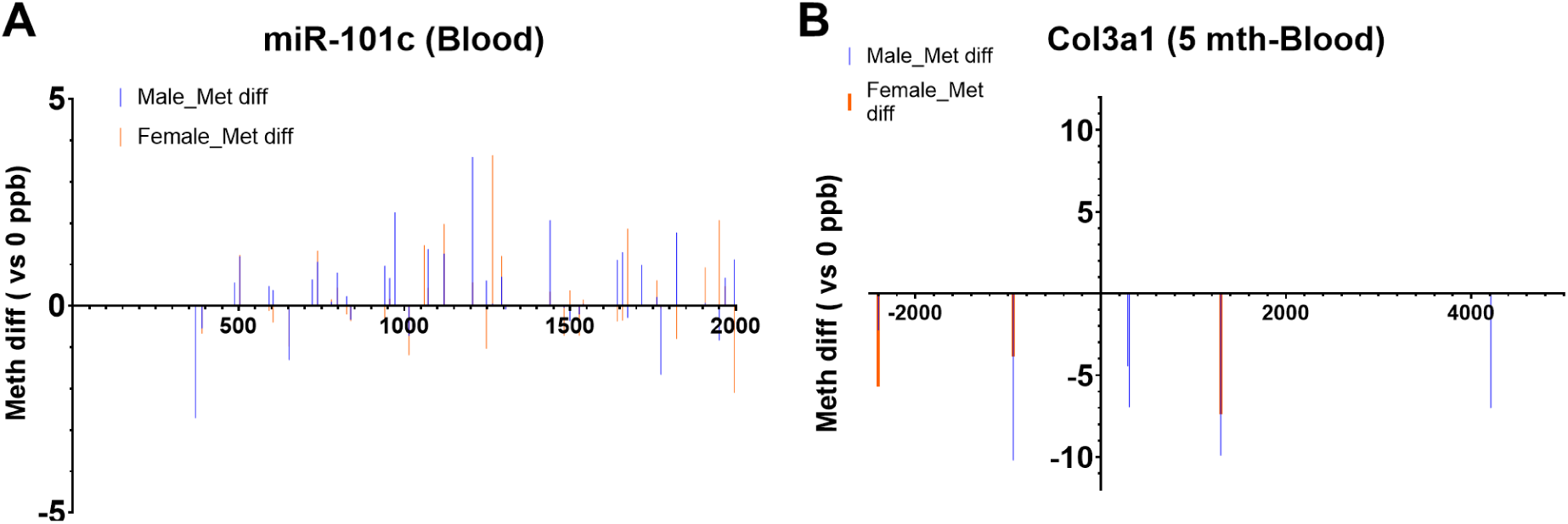
Methylation differences (from WGBS) in *miR-101c* and *col3a1* between controls and iAs-exposed group in male and female blood leukocytes. Differences were identified using the methylKit package, selecting CpGs with a minimum read depth of 10X across samples and a q-value ≤ 0.05.

**Supplementary Table 1.**
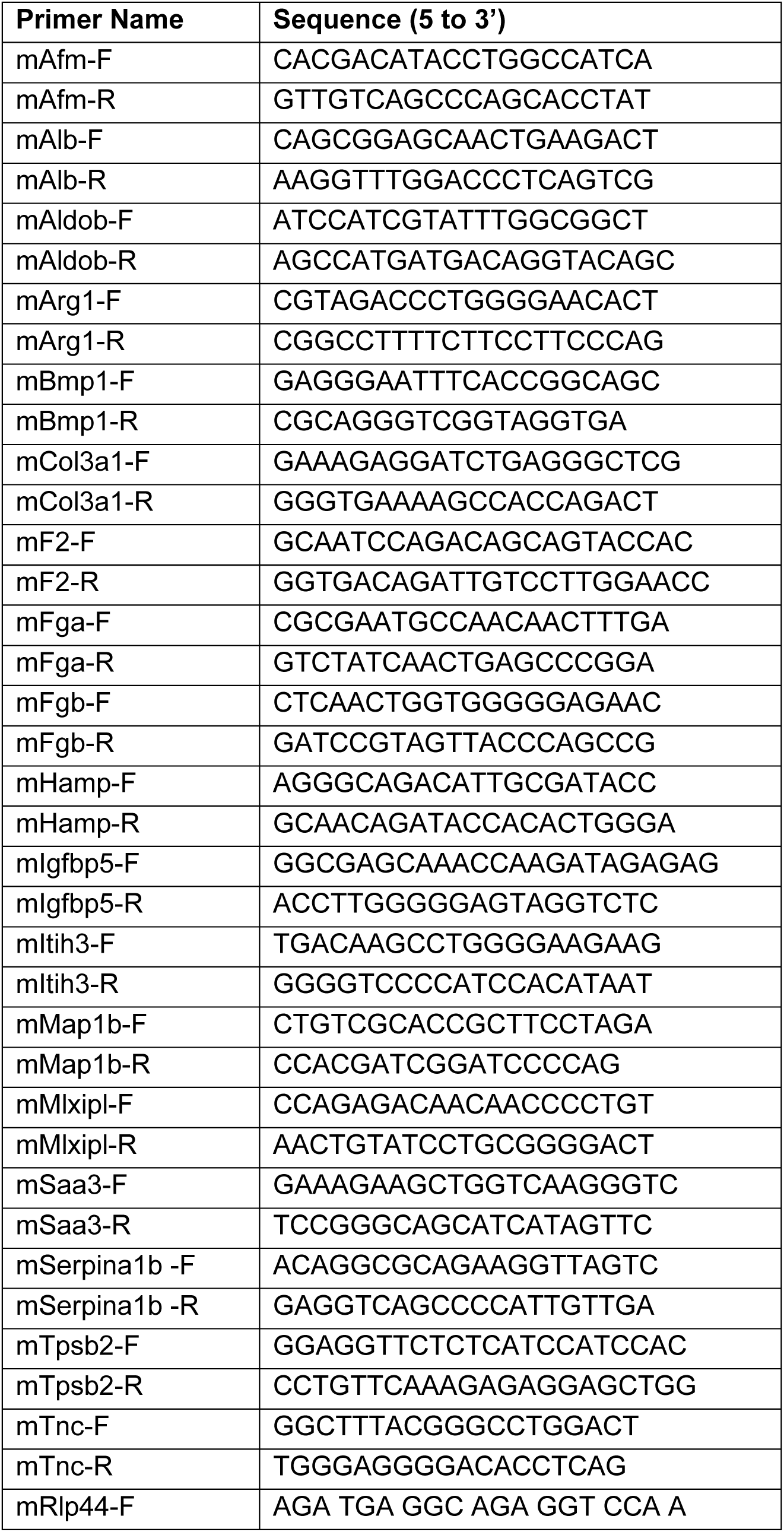
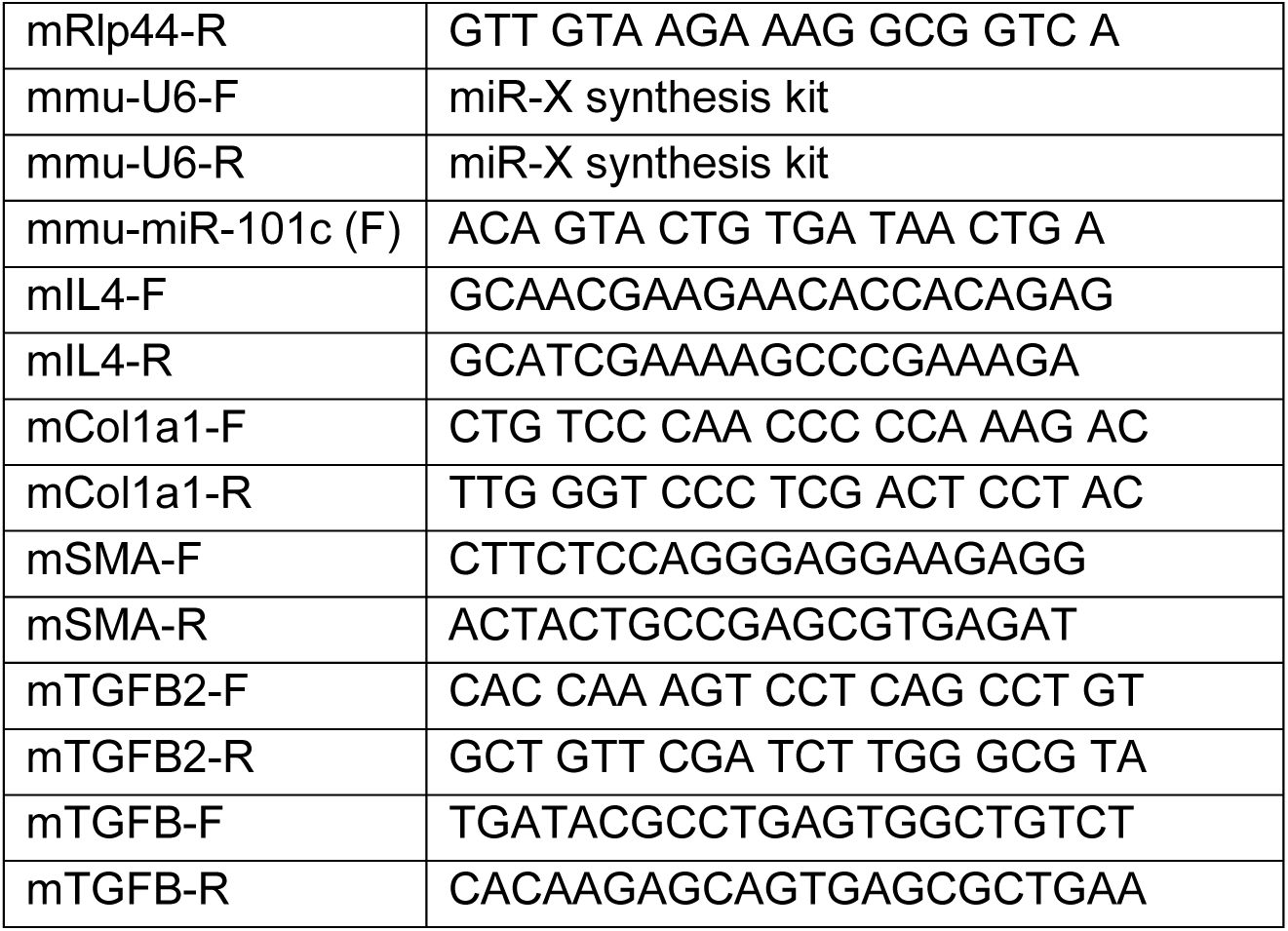
Primer sequences for qRT-PCR.

**Supplementary Table 2.** List of seventy differential expressed genes (DEGs) common in male and female blood samples. Seven of them are not RefSeq genes.

| Gene symbol | Entrezgene | Gene Name | taxid | MGI |
| --- | --- | --- | --- | --- |
| abcc3 | 76408 | ATP-binding cassette, sub-family C (CFTR/MRP), member 3 | 10090 | MGI:1923658 |
| ac159819.1 | n.a. | n.a. | n.a. | n.a. |
| acaa1b | 235674 | acetyl-Coenzyme A acyltransferase 1B | 10090 | MGI:3605455 |
| afm | 280662 | afamin | 10090 | MGI:2429409 |
| agxt | 11611 | alanine-glyoxylate aminotransferase | 10090 | MGI:1329033 |
| alb | 11657 | albumin | 10090 | MGI:87991 |
| aldob | 230163 | aldolase B, fructose-bisphosphate | 10090 | MGI:87995 |
| ankle1 | 234396 | ankyrin repeat and LEM domain containing 1 | 10090 | MGI:1918775 |
| apoa4 | 11808 | apolipoprotein A-IV | 10090 | MGI:88051 |
| arg1 | 94242 | tubulointerstitial nephritis antigen-like 1 | 10090 | MGI:2137617 |
| atat1 | 73242 | alpha tubulin acetyltransferase 1 | 10090 | MGI:1913869 |
| baiap2l1 | 66898 | BAI1-associated protein 2-like 1 | 10090 | MGI:1914148 |
| bhmt | 12116 | betaine-homocysteine methyltransferase | 10090 | MGI:1339972 |
| bmp1 | 12153 | bone morphogenetic protein 1 | 10090 | MGI:88176 |
| c77080 | 97130 | expressed sequence C77080 | 10090 | MGI:2140651 |
| cchcr1 | 240084 | coiled-coil alpha-helical rod protein 1 | 10090 | MGI:2385321 |
| ces1c | 13884 | carboxylesterase 1C | 10090 | MGI:95420 |
| col3a1 | 12825 | collagen, type III, alpha 1 | 10090 | MGI:88453 |
| cyp3a25 | 56388 | cytochrome P450, family 3, subfamily a, polypeptide 25 | 10090 | MGI:1930638 |
| cyp4a10 | 13117 | cytochrome P450, family 4, subfamily a, polypeptide 10 | 10090 | MGI:88611 |
| cyp4a14 | 13119 | cytochrome P450, family 4, subfamily a, polypeptide 14 | 10090 | MGI:1096550 |
| cyp8b1 | 13124 | cytochrome P450, family 8, subfamily b, polypeptide 1 | 10090 | MGI:1338044 |
| ephx2 | 13850 | epoxide hydrolase 2, cytoplasmic | 10090 | MGI:99500 |
| espn | 56226 | espin | 10090 | MGI:1861630 |
| f2 | 14061 | coagulation factor II | 10090 | MGI:88380 |
| fabp1 | 14080 | fatty acid binding protein 1, liver | 10090 | MGI:95479 |
| fga | 14161 | fibrinogen alpha chain | 10090 | MGI:1316726 |
| fgb | 110135 | fibrinogen beta chain | 10090 | MGI:99501 |
| ftcd | 14317 | formiminotransferase<br>cyclodeaminase | 10090 | MGI:1339962 |
| gm10717 | n.a. | n.a. | n.a. | n.a. |
| gm28035 | n.a. | n.a. | n.a. | n.a. |
| gm28539 | n.a. | n.a. | n.a. | n.a. |
| gm38431 | n.a. | n.a. | n.a. | n.a. |
| gm49345 | n.a. | n.a. | n.a. | n.a. |
| gm49358 | n.a. | n.a. | n.a. | n.a. |
| hamp | 84506 | hepcidin antimicrobial peptide | 10090 | MGI:1933533 |
| hmgcs2 | 15360 | 3-hydroxy-3-methylglutaryl-Coenzyme A synthase 2 | 10090 | MGI:101939 |
| htra3 | 78558 | HtrA serine peptidase 3 | 10090 | MGI:1925808 |
| igfbp5 | 16011 | insulin-like growth factor binding<br>protein 5 | 10090 | MGI:96440 |
| itih2 | 16425 | inter-alpha trypsin inhibitor,<br>heavy chain 2 | 10090 | MGI:96619 |
| itih3 | 16426 | inter-alpha trypsin inhibitor,<br>heavy chain 3 | 10090 | MGI:96620 |
| map1b | 17755 | microtubule-associated protein<br>1B | 10090 | MGI:1306778 |
| mat1a | 11720 | methionine adenosyltransferase<br>I, alpha | 10090 | MGI:88017 |
| mlxipl | 58805 | MLX interacting protein-like | 10090 | MGI:1927999 |
| mup11 | 100039028 | major urinary protein 11 | 10090 | MGI:3709617 |
| mup18 | 100048884 | major urinary protein 18 | 10090 | MGI:3705220 |
| nlgn3 | 245537 | neuroligin 3 | 10090 | MGI:2444609 |
| nr1i3 | 12355 | nuclear receptor subfamily 1,<br>group I, member 3 | 10090 | MGI:1346307 |
| pck1 | 18534 | phosphoenolpyruvate<br>carboxykinase 1, cytosolic | 10090 | MGI:97501 |
| phldb1 | 102693 | pleckstrin homology like<br>domain, family B, member 1 | 10090 | MGI:2143230 |
| prg2 | 216152 | phospholipid phosphatase<br>related 3 | 10090 | MGI:2388640 |
| proc | 19123 | protein C | 10090 | MGI:97771 |
| prodh2 | 56189 | proline dehydrogenase<br>(oxidase) 2 | 10090 | MGI:1929093 |
| proz | 66901 | protein Z, vitamin K-dependent<br>plasma glycoprotein | 10090 | MGI:1860488 |
| prrt2 | 69017 | proline-rich transmembrane<br>protein 2 | 10090 | MGI:1916267 |
| rps18-ps6 | 100039740 | ribosomal protein S18,<br>pseudogene 6 | 10090 | MGI:3642298 |
| saa3 | 20210 | serum amyloid A 3 | 10090 | MGI:98223 |
| sds | 66711 | SBDS ribosome maturation<br>factor | 10090 | MGI:1913961 |
| sec14l2 | 67815 | SEC14-like lipid binding 2 | 10090 | MGI:1915065 |
| serpina1b | 20701 | serine (or cysteine) preptidase<br>inhibitor, clade A, member 1B | 10090 | MGI:891970 |
| serpinc1 | 11905 | serine (or cysteine) peptidase inhibitor, clade C (antithrombin), member 1 | 10090 | MGI:88095 |
| slc12a5 | 57138 | solute carrier family 12, member 5 | 10090 | MGI:1862037 |
| sorbs3 | 20410 | sorbin and SH3 domain containing 3 | 10090 | MGI:700013 |
| tat | 234724 | tyrosine aminotransferase | 10090 | MGI:98487 |
| tdo2 | 56720 | tryptophan 2,3-dioxygenase | 10090 | MGI:1928486 |
| tnc | 21924 | troponin C, cardiac/slow skeletal | 10090 | MGI:98779 |
| tpsb2 | 17229 | tryptase beta 2 | 10090 | MGI:96942 |
| tub | 22141 | tubby bipartite transcription factor | 10090 | MGI:2651573 |
| uroc1 | 243537 | urocanase domain containing 1 | 10090 | MGI:2385332 |
| zfp811 | 240063 | zinc finger protein 811 | 10090 | MGI:2682944 |

**Supplementary Table 3.**
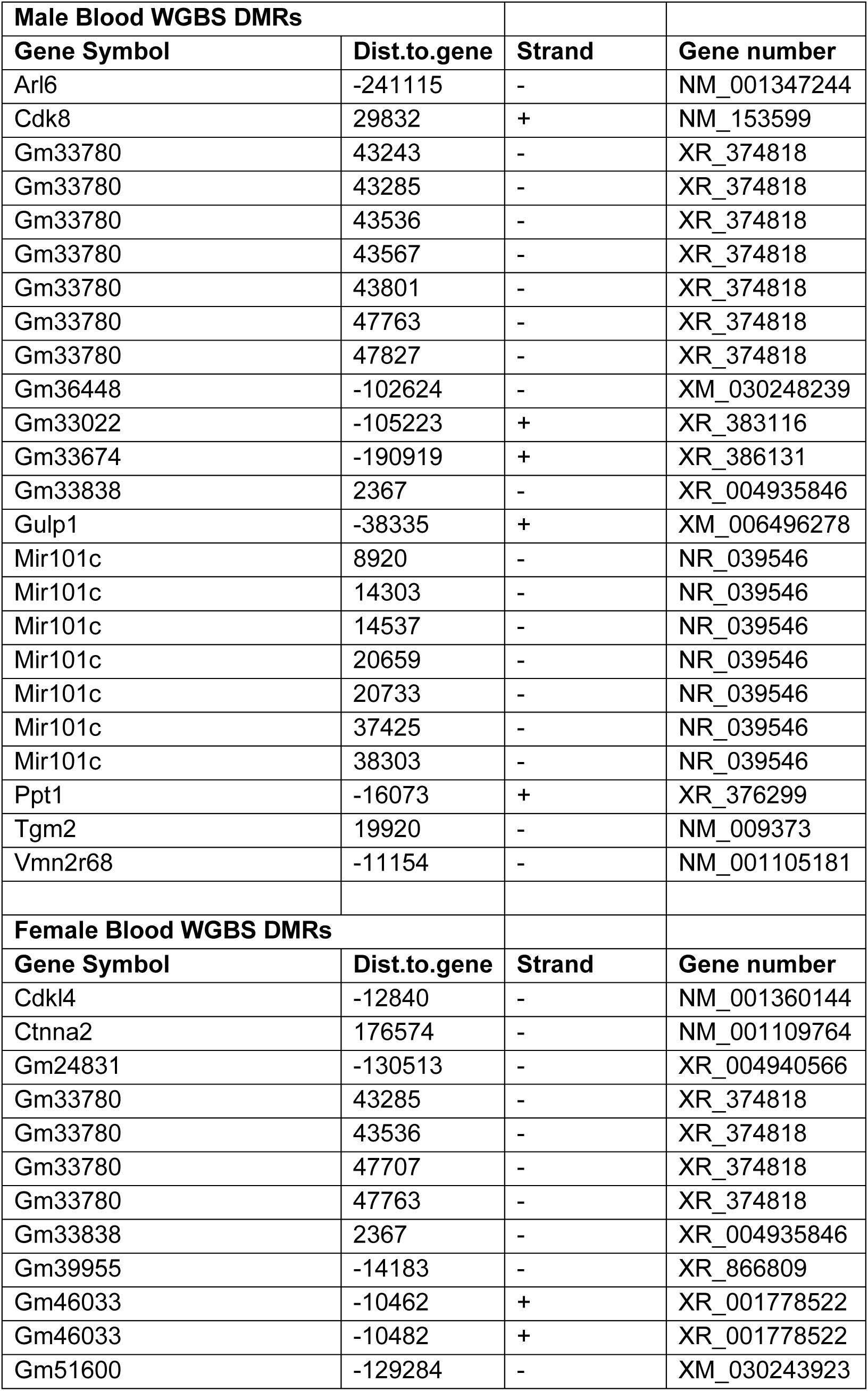
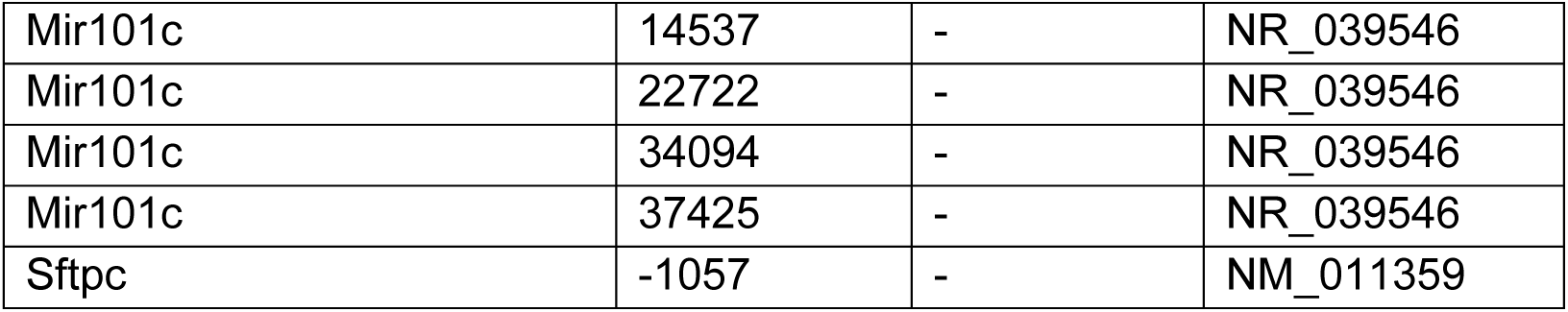
List of DMCs in male and female blood samples.

**Supplementary Table 4.** Expression of DEGs (common in male and female) in surrogate (blood) and target (lung) tissues. Orange: ECM signaling-related genes; yellow: inflammatory response-related genes; blue: embryonic development-related genes. ^ Indicates differentially methylated candidates.

| Male<br>Gene<br>Symbol | 3-week-old<br>(Blood<br>Leukocytes) |  | 3-week-old<br>(Lung) |  | 5-mth-old (Blood<br>Leukocytes) |  | 5-month-old<br>(Lung) |  |
| --- | --- | --- | --- | --- | --- | --- | --- | --- |
|  | Log2<br>FC | p value | Log2<br>FC | p value | Log2<br>FC | p value | Log2<br>FC | p value |
| Arg1 | 0.710 | 0.560 | -0.073 | 0.867 | 5.620 | **<0.001** | 0.254 | 0.169 |
| Bmp1 | -1.430 | 0.140 | -0.102 | 0.741 | 5.450 | **0.010** | 0.287 | 0.567 |
| Col3a1 | -0.580 | 0.430 | 0.837 | **0.015** | 3.930 | **0.020** | 0.700 | **0.036** |
| Itih3 | 6.530 | **0.004** | -1.132 | **0.032** | 3.830 | **0.001** | -0.678 | **0.024** |
| Map1b | -2.940 | **0.002** | 0.639 | 0.285 | 3.450 | **0.009** | 0.851 | **0.019** |
| Tnc | 2.460 | **0.050** | 0.175 | 0.742 | 5.510 | **0.020** | 0.807 | 0.147 |
| Tpsb2 | -2.030 | 0.060 | 0.938 | **0.005** | 6.100 | **0.005** | 1.937 | **0.019** |
| Fga | 7.190 | **0.002** | 1.839 | **0.025** | 2.520 | **0.005** | -0.673 | 0.072 |
| Fgb | 2.820 | **0.020** | 2.350 | **0.042** | 4.080 | **<0.001** | 4.419 | **0.050** |
| F2 | 3.750 | **0.010** | 0.606 | 0.409 | 6.100 | **<0.001** | 2.039 | **0.015** |
| Hamp | 2.630 | 0.230 | -0.088 | 0.843 | 3.640 | **0.001** | -1.681 | **0.027** |
| MIxipl | -2.250 | **0.040** | 0.569 | 0.200 | 4.470 | **<0.001** | 0.342 | 0.459 |
| Saa3 | -2.230 | **0.001** | -0.164 | 0.683 | 2.750 | **0.004** | 1.283 | **0.018** |
| Afm | 1.990 | 0.210 | 0.432 | 0.548 | 4.960 | **0.030** | 0.284 | 0.807 |
| Alb | 5.400 | **<0.001** | 1.330 | 0.194 | 3.520 | **<0.001** | 1.010 | 0.100 |
| Aldob | 3.360 | **0.003** | 2.294 | 0.091 | 3.570 | **<0.001** | 1.334 | **0.028** |
| Igfbp5 | -1.850 | **0.002** | 0.415 | 0.207 | 2.280 | **0.010** | 0.693 | 0.239 |
| Serpina1b | 4.920 | **<0.001** | 1.766 | 0.230 | 3.080 | **0.002** | 1.551 | **0.015** |
| ^miR-101c | -1.270 | **0.039** | -0.536 | 0.190 | -2.535 | **0.048** | -1.706 | **0.044** |

| Female<br>Gene<br>Symbol | 3-week-old<br>(Blood<br>Leukocytes) |  | 3-week-old<br>(Lung) |  | 5-mth-old (Blood<br>Leukocytes) |  | 5-month-old<br>(Lung) |  |
| --- | --- | --- | --- | --- | --- | --- | --- | --- |
|  | Log2<br>FC | p value | Log2<br>FC | p value | Log2<br>FC | p value | Log2<br>FC | p value |
| Arg1 | 1.610 | 0.150 | -0.188 | 0.661 | -3.150 | **0.002** | -0.537 | 0.253 |
| Bmp1 | 2.790 | **0.005** | 0.407 | 0.155 | -2.190 | **0.010** | -0.174 | 0.562 |
| Col3a1 | 3.350 | **<0.001** | 1.392 | **0.008** | 3.250 | **0.001** | 0.538 | **0.021** |
| Itih3 | -1.560 | 0.290 | -1.216 | 0.024 | -2.450 | **0.030** | -0.494 | **0.019** |
| Map1b | 1.150 | 0.260 | 0.543 | 0.121 | -3.930 | **<0.001** | 0.372 | 0.210 |
| Tnc | 4.580 | **0.001** | 0.439 | 0.399 | -2.740 | **0.030** | -0.133 | 0.680 |
| Tpsb2 | 2.630 | **0.020** | 1.548 | **0.048** | -4.920 | **<0.001** | 1.133 | **0.021** |
| Fga | -1.810 | 0.200 | 1.853 | **0.054** | -1.950 | **0.050** | 0.204 | 0.626 |
| Fgb | -0.360 | 0.790 | 2.970 | **0.019** | -2.800 | **0.003** | 3.498 | **0.043** |
| F2 | -1.170 | 0.380 | 0.359 | 0.294 | -2.470 | **0.010** | -0.006 | 0.985 |
| Hamp | ND | ND | -0.602 | **0.024** | -2.350 | **0.030** | -1.228 | **0.040** |
| Mlxipl | 3.410 | **0.001** | 0.647 | 0.153 | -3.200 | **0.002** | -0.120 | 0.697 |
| Saa3 | 3.200 | **0.001** | 1.079 | **0.029** | -2.210 | **0.020** | 1.289 | **0.029** |
| Afm | 1.750 | 0.260 | 1.193 | **0.008** | -3.900 | **0.010** | 0.740 | **0.044** |
| Alb | 0.030 | 0.970 | 0.951 | 0.180 | -2.150 | **0.005** | 0.130 | 0.833 |
| Aldob | -0.980 | 0.430 | 1.171 | **0.049** | -2.200 | **0.010** | 0.641 | **0.040** |
| Igfbp5 | 0.320 | 0.610 | 0.592 | 0.114 | 3.320 | **<0.001** | 0.077 | 0.798 |
| Serpina1b | -0.650 | 0.600 | -0.292 | 0.576 | -1.800 | **0.030** | 2.983 | 0.051 |
| ^miR-101c | -2.024 | **0.007** | -0.828 | **0.042** | -0.782 | **0.039** | -1.904 | **0.009** |

**Supplementary Table 5:**
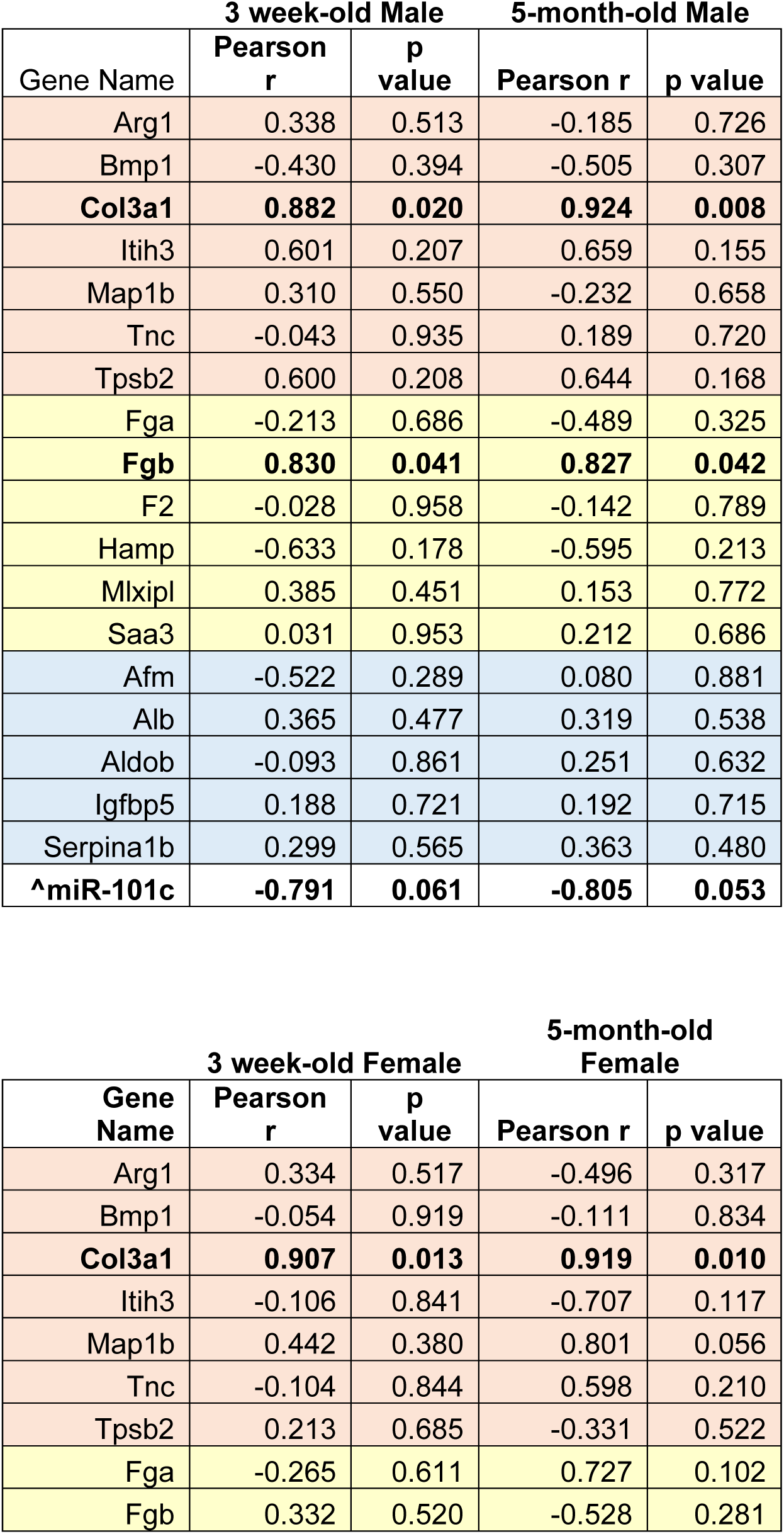
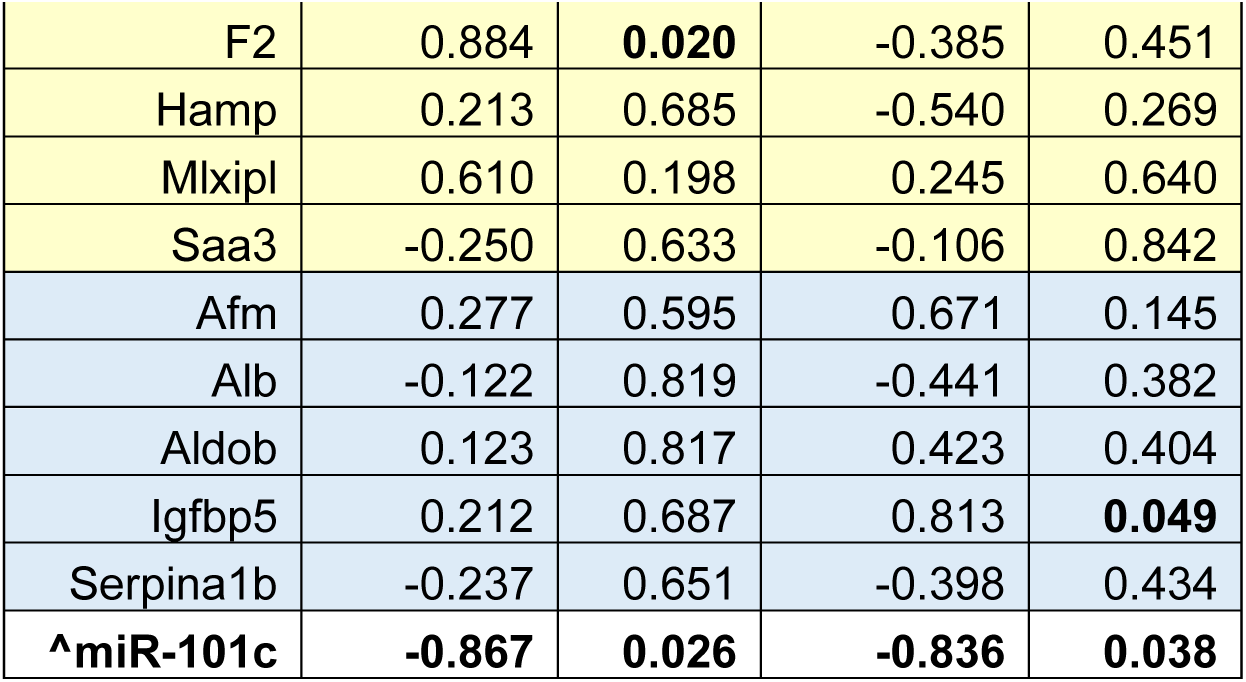
Correlation analysis of iAs-induced blood transcriptional changes and asthma risk. Orange: ECM signaling-related genes; yellow: inflammatory response-related genes; blue: embryonic development-related genes. ^ Indicates differentially methylated candidates.

|  | 3 week-old Male |  | 5-month-old Male |  |
| --- | --- | --- | --- | --- |
| Gene Name | Pearson r | p value | Pearson r | p value |
| Arg1 | 0.338 | 0.513 | -0.185 | 0.726 |
| Bmp1 | -0.430 | 0.394 | -0.505 | 0.307 |
| **Col3a1** | **0.882** | **0.020** | **0.924** | **0.008** |
| Itih3 | 0.601 | 0.207 | 0.659 | 0.155 |
| Map1b | 0.310 | 0.550 | -0.232 | 0.658 |
| Tnc | -0.043 | 0.935 | 0.189 | 0.720 |
| Tpsb2 | 0.600 | 0.208 | 0.644 | 0.168 |
| Fga | -0.213 | 0.686 | -0.489 | 0.325 |
| **Fgb** | **0.830** | **0.041** | **0.827** | **0.042** |
| F2 | -0.028 | 0.958 | -0.142 | 0.789 |
| Hamp | -0.633 | 0.178 | -0.595 | 0.213 |
| Mlxipl | 0.385 | 0.451 | 0.153 | 0.772 |
| Saa3 | 0.031 | 0.953 | 0.212 | 0.686 |
| Afm | -0.522 | 0.289 | 0.080 | 0.881 |
| Alb | 0.365 | 0.477 | 0.319 | 0.538 |
| Aldob | -0.093 | 0.861 | 0.251 | 0.632 |
| Igfbp5 | 0.188 | 0.721 | 0.192 | 0.715 |
| Serpina1b | 0.299 | 0.565 | 0.363 | 0.480 |
| **^miR-101c** | **-0.791** | **0.061** | **-0.805** | **0.053** |

|  | 3 week-old Female |  | 5-month-old Female |  |
| --- | --- | --- | --- | --- |
| Gene Name | Pearson r | p value | Pearson r | p value |
| Arg1 | 0.334 | 0.517 | -0.496 | 0.317 |
| Bmp1 | -0.054 | 0.919 | -0.111 | 0.834 |
| **Col3a1** | **0.907** | **0.013** | **0.919** | **0.010** |
| Itih3 | -0.106 | 0.841 | -0.707 | 0.117 |
| Map1b | 0.442 | 0.380 | 0.801 | 0.056 |
| Tnc | -0.104 | 0.844 | 0.598 | 0.210 |
| Tpsb2 | 0.213 | 0.685 | -0.331 | 0.522 |
| Fga | -0.265 | 0.611 | 0.727 | 0.102 |
| Fgb | 0.332 | 0.520 | -0.528 | 0.281 |
| F2 | 0.884 | **0.020** | -0.385 | 0.451 |
| Hamp | 0.213 | 0.685 | -0.540 | 0.269 |
| Mlxip1 | 0.610 | 0.198 | 0.245 | 0.640 |
| Saa3 | -0.250 | 0.633 | -0.106 | 0.842 |
| Afm | 0.277 | 0.595 | 0.671 | 0.145 |
| Alb | -0.122 | 0.819 | -0.441 | 0.382 |
| Aldob | 0.123 | 0.817 | 0.423 | 0.404 |
| Igfbp5 | 0.212 | 0.687 | 0.813 | **0.049** |
| Serpina1b | -0.237 | 0.651 | -0.398 | 0.434 |
| **^miR-101c** | **-0.867** | **0.026** | **-0.836** | **0.038** |

## DATA AVAILABILITY

RNA-seq and WGBS data in the paper has been deposited through Gene Expression Omnibus (GEO) repository: GSE146508 and those data are also available at the TaRGET II data portal (https://data.targetepigenomics.org/).

## Notes

### Competing Interest Statement

The authors have declared no competing interest.

